# HyDrop v2: Scalable atlas construction for training sequence-to-function models

**DOI:** 10.1101/2025.04.02.646792

**Authors:** Hannah Dickmänken, Marta Wojno, Koen Theunis, Eren Can Ekşi, Lukas Mahieu, Valerie Christiaens, Niklas Kempynck, Florian V. De Rop, Natalie Roels, Katina I. Spanier, Roel Vandepoel, Gert Hulselmans, Suresh Poovathingal, Stein Aerts

## Abstract

Deciphering cis-regulatory logic underlying cell type identity is a fundamental question in biology. Single-cell chromatin accessibility (scATAC-seq) data has enabled training of sequence-to-function deep learning models allowing decoding of enhancer logic and design of synthetic enhancers. Training such models requires large amounts of high-quality training data across species, organs, development, aging, and disease. To facilitate the cost-effective generation of large scATAC-seq atlases for model training, we developed a new version of the open-source microfluidic system HyDrop with increased sensitivity and scale: HyDrop v2. We generated HyDrop v2 atlases for the mouse cortex and *Drosophila* embryo development and compared them to atlases generated on commercial platforms. HyDrop v2 data integrates seamlessly with commercially available chromatin accessibility methods (10x Genomics). Differentially accessible regions and motif enrichment across cell types are equivalent between HyDrop-v2 and 10x atlases. Sequence-to-function models trained on either atlas are comparable as well in terms of enhancer predictions, sequence explainability, and transcription factor footprinting. By offering accessible data generation, enhancer models trained on HyDrop-v2 and mixed atlases can contribute to unraveling cell-type specific regulatory elements in health and disease.

## Introduction

Cell type identity is encoded in the DNA sequence of *cis*-regulatory elements (CREs), such as enhancers^1^. By harboring binding sites for specific combinations of transcription factors (TFs), CREs orchestrate cell type-specific binding of TFs subsequently resulting in the expression of genes underlying the cell’s identity. Thus, identifying CREs is a fundamental question in biology. Several experimental approaches are available to identify CREs and characterize their function^1,2^. Two commonly used approaches are (1) genome-wide profiling of TF binding, with assays such as Chromatin Immuno Precipitation sequencing (ChIP-seq)^3^, Cleavage Under Target and Release Using Nuclease (CUT& RUN)^4^, and Cleavage Under Targets and Tagmentation (CUT&Tag)^2^; and (2) genome-wide profiling of chromatin activity such as histone modifications and chromatin accessibility^1^. Measuring chromatin accessibility with Assay of Transposase Accessible Chromatin (ATAC) offers the most unbiased approach that can be performed in single cells (scATAC)^1^.

Various tools are available to process scATAC data and identify cell-type-specific regulatory sequences^5–9^ (i.e., differentially accessible regions, DARs). Combining scATAC with gene expression information, regulatory regions can be linked to downstream target gene expression^10,11^. The rapid growth of machine learning, especially deep learning (DL) has opened new possibilities for accurately identifying specific TF binding sites (TFBS) within accessible regions. So-called sequence-to-function (S2F) models identify the nucleotide sequence patterns of regulatory elements in the non-coding genome with only scATAC information as training data^12–15^ (for an in-depth review see De Winter et al. (2025)^16^).

However, the combination of the large demand for training data for DL models^12,15^ and the sparse nature of scATAC data^1^ results in a pressing need for large-scale data sets across species, genotypes, conditions, and developmental stages. Overcoming the challenge of limited scATAC enhancer data is possible by DL models using orthogonal data, as shown previously in work on the fly embryo^15^. This, however, requires additional information from large-scale *in vivo* experiments^15,17^ that are not always readily available. In contrast, other groups have shown that training high-quality S2F models with merely scATAC data is possible in fly^14^ and other (model) organisms^12–14^. Along the same line, a recent benchmarking study on various data modalities showed that training DL models on scATAC data only perform superior in predicting cell type-specific enhancers than non-DL methods or using other data modalities^12^. Importantly, the need for high-quality scATAC data with improved coverage is emphasized^12^.

Most large-scale data sets focus on mouse data or cell lines, while less benchmarking is done on non-vertebrate model organisms, leading to an underrepresentation in the data landscape^12,13,18^. However, including complex tissues and organisms with more than a handful of cell types is essential as the computational identification of functional elements paves the way for fundamental biological questions on cell state regulation. Typically, experimental validation is costly and difficult to leverage on a large scale, especially for complex model organisms compared to cell lines. Enhancer prediction with S2F models enhances speed and accuracy towards wet lab experiments to eventually understand the grammar of cell-type specific regulatory sequences^12,14,15^.

Over the recent years, various scATAC-seq methods have been developed primarily using either well-based combinatorial indexing strategies or microfluidics encapsulation technology (for an in-depth comparison, see De Rop et al., 2023^19^). In an earlier publication, we developed HyDrop, a microfluidics droplet technology based on deformable hydrogel indexing beads, which enables high-throughput profiling of both single-cell RNAseq and single nuclei ATAC seq^20^. Based on this work, we have improved several aspects of the technology to develop HyDrop-ATAC v2 (further addressed as HyDrop v2), which enables profiling of single-cell ATAC seq with higher detection sensitivity. Like commercial platforms for droplet-based scATAC-seq (e.g., 10X Genomics Epi ATAC), HyDrop v2 uses a hydrogel-based single-cell indexing strategy with droplet microfluidics. Alternative commercial platforms are based on microwell encapsulation of single cells (BD Rhapsody) or use a well-based combinatorial indexing strategy (Scale Bioscience). Overall, these commercial platforms are costly and/or require labor-intensive workflows that hamper the generation of large-scale single-cell atlasing works of complex tissues^19^. Importantly, beyond comparing the quality and costs of various scATAC techniques^19^, no benchmarking has been performed to assess the effect of different scATAC-seq techniques (e.g., commercial versus custom platforms) on their capacity to train S2F models. Such a quantitative evaluation of scATAC techniques becomes even more relevant in light of the facilitated use of DL for a broad research audience^6,21,22^.

To address the need for high-quality and low-cost scATAC data, we refined the open-source microfluidic platform HyDrop. The next version, HyDrop v2, offers improved and consistent data quality across multiple batches of bead production. In cutting-edge S2F models and footprinting predictions in mouse cortex and fly embryo, HyDrop v2 data perform analogously to commercial data. By offering highly accessible data generation with the open-source HyDrop v2, we contribute to unraveling cell-type specific regulatory elements in health and disease with deep learning.

## Results

### HyDrop bead chemistry optimization enables reproducible fragment capturing

The bead barcoding chemistry in the previous version of HyDrop^20^ was performed using a 3-stage extension of DNA primers grafted onto polyacrylamide gel beads that were generated using droplet microfluidics. This was achieved with 96 different barcoded DNA primers. Three-stage barcoding with 96 barcoded DNA primers results in a barcode complexity of 96^3^ = 884,736 barcodes. During each stage of this barcoding strategy, hydrogel beads were mixed with DNA polymerase and a uniquely barcoded extension primer (Figure 1a). The extension of the grafted primers in the beads was achieved by cyclic DNA polymerase extension. Before the next stage of bead barcoding, the second strand was denatured, and beads were washed to remove the second strand (Figure 1a). In the current version of HyDrop v2 bead production, the polymerase-based cyclic extension of the barcoded primers was replaced by ligation of the barcoded primers. Barcoded oligo cassettes are now ligated sequentially in a 3-stage barcoding process resulting in same barcode complexity as the HyDrop v1 beads. Importantly, in the HyDrop v2 bead production, there is no intermediate strand denaturation or cyclic addition of primers to beads (Figure 1a), thereby enhancing the barcode stability (Figure 1d).

**Fig. 1.**
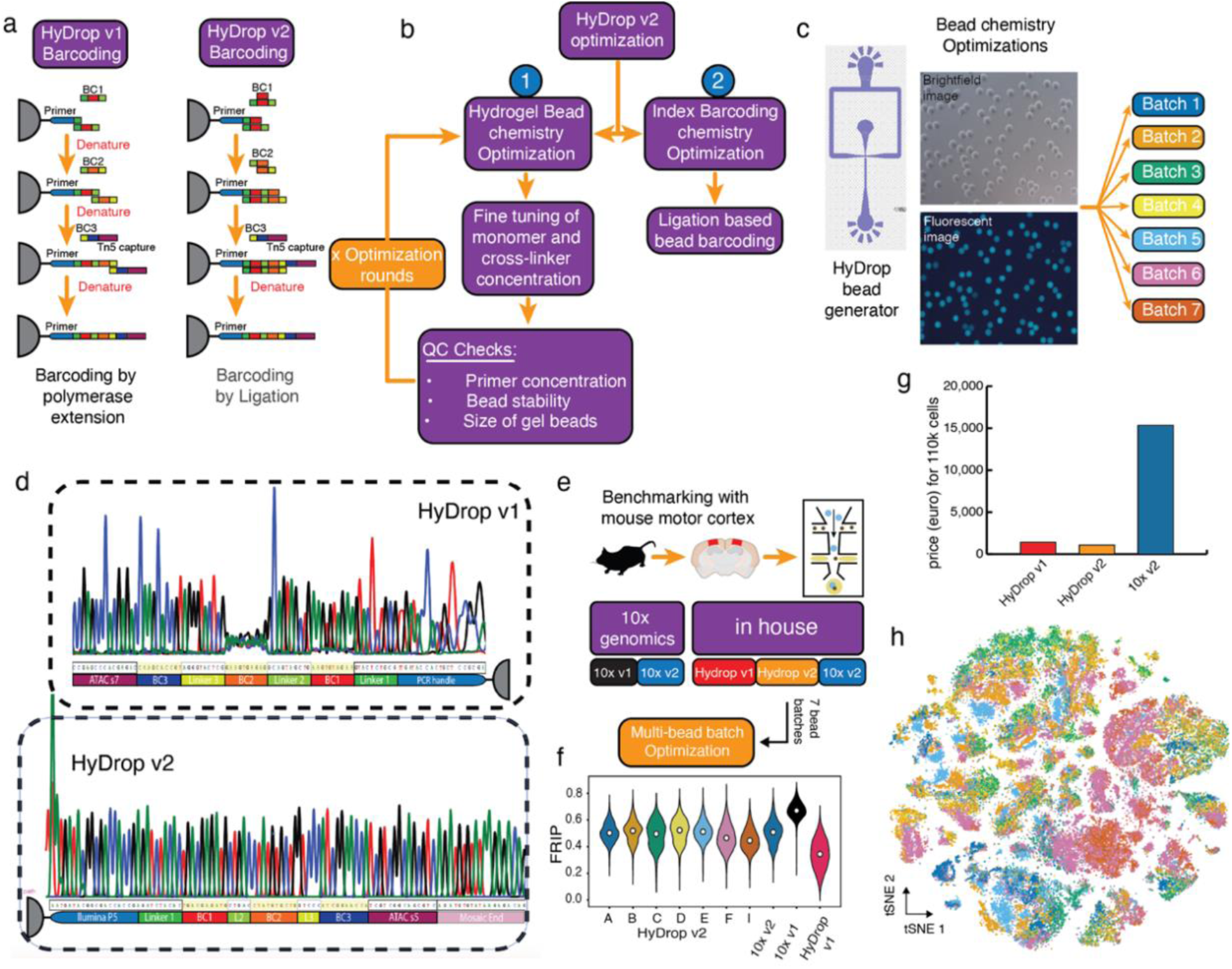
Optimizations of HyDrop enable consistently high fragment capture. **a**, Comparison between bead barcoding chemistries of HyDrop v1 and v2. **b**, The HyDrop v1 protocol was optimized in two ways: (1) Bead chemistry was adjusted in several optimization rounds, and (2) the barcoding of the beads was changed from a cyclic polymerase extension to a linear ligation-based chemistry. **c**, The design of the microfluidic chip to generate HyDrop hydrogel beads (left), alongside the image of the polymerized hydrogel bead and fluorescence image of the bead labeled with complement of Tn5 fragment capture site. **d,** Sanger sequencing of an exemplary HyDrop v1 bead (top) showing impurity in barcode one and barcode two and a HyDrop v2 bead with pure barcode signals across all three barcodes. **e**, Mouse cortex data were partly downloaded from the official 10x genomics website and partly generated in-house. Overview of the sample generation: dissection of mouse motor cortex, tissue lysis followed by droplet encapsulation for cell barcoding. For HyDrop v1 only one bead batch was generated (data reused from De Rop et al., 2022) while for HyDrop v2 seven bead batches were generated (seen in c). **f**, FRIP of experiments pooled for seven different bead batches of HyDrop v2 compared to pooled 10x v2, 10x v1, and HyDrop v1 samples. The median is indicated per sample in white. **g,** Estimation of generation costs of 110k cells in euros excluding sequencing costs. HyDrop v1: 1,398.29 euros, HyDrop v2: 1,080.60 euros, 10x v2: 15,330.70 euros. **h**, *t*-distributed stochastic neighbor embedding (tSNE) of all 110,547 cells generated with HyDrop v2 (35 experiments) colored by bead batch (seen in c), no batch correction. Data points were randomly shuffled before plotting. QC: Quality control, FRIP: Fraction of reads in peaks.

We compared the purity of individual barcodes per bead by single bead picking using a micro-injection syringe followed by amplification and Sanger sequencing (Fig 1d, Fig S1f). Compared to HyDrop v1, which showed higher barcode contamination across different barcodes, all three ligated barcodes in the v2 beads are correctly individualized per bead.

An additional quality challenge that we faced with HyDrop v1 bead production was the maintenance of stable bead quality, both in terms of the stability of the polyacrylamide gel bead matrix as well as in the maintenance of the size of the gel beads during bead production. Both factors are closely tied to the monomer composition of polyacrylamide chemistry. Earlier works have demonstrated the structural stability of polyacrylamide chemistry is governed by the monomer and crosslinker compositions^23^. Accordingly, a series of optimization cycles improving the composition of total monomer concentration (%T) and crosslinker concentration (%C) (Figure S1a) was performed. Based on different conditions we found 10% total monomer concentration (T) and 2% crosslinker concentration to produce the most stable beads (Figure S1a). These changes in bead generation were validated through multiple batches of beads that were produced across seven independent bead production runs (Figure 1C, Figure S1d). The bead batches were generated using an in-house fabricated microfluidics droplet generator as described earlier^20^. A quality comparison of the hydrogel beads produced with commercial alternates (droplet generator chips from Atrandi Biosciences) showed that the gel beads generated with the in-house fabricated bead generator yield beads of superior quality (Figure S1d, e). This is primarily attributed to the incompatibility of adding the N,N,N′,N′ -Tetramethylethylenediamine (TEMED), a polyacrylamide polymerization catalyst with the Atrandi’s droplet generation chips. TEMED was added post droplet generation to the bead emulsion oil. Droplets generated using commercial chips furthermore result in freezing artifacts from bead storage (Figure S1e).

We evaluated the changes in bead chemistry and generation by comparing commercial microfluidics platforms from 10x genomics to HyDrop v1 (one bead batch) and HyDrop v2 using mouse motor cortex tissue (Figure 1e). For HyDrop v2, seven independent bead batches were used, showing high data integrability without the need for batch correction (Figure 1h). The described changes in HyDrop v2 compared to HyDrop v1 result in improved performance in recovered unique fragments with an increase of 10.06%. Here, HyDrop v2 reaches, on average, 93.01% (M HyDrop v2=3.92, SD=0.38; M 10x v2=4.21, SD=0.27) of the unique number of fragments (log scaled) captured with 10x v2 (Figure S1b). Similarly, the fraction of reads found in peaks (FRIP, Figure 1f) in HyDrop v2 and the sequencing efficiency (Figure S2b) of Hydrop v2 nearly reached 10x genomics levels (M FRIP HyDrop v2=0.49, SD=0.09; M FRIP 10x v2=0.51, SD=0.09).

We generated an atlas of 110k cells covering the major cell types in the mouse motor cortex using the HyDrop v2 platform for 1,080.6 euros (excluding sequencing). Generating a data set of this size using HyDrop v1 would have cost 1,398.29 euros while using the current gold standard of scATAC sequencing (10x v2 chemistry) would result in a cost of approximately 15,330.70 euros (Figure 1g).

Here, we demonstrate that the improvements from HyDrop v1 to v2 chemistry result in reliable data generation of large-scale scATAC data sets in complex tissue covering various cell types. Additionally, we were able to reduce the costs of HyDrop even further while increasing the sequencing efficiency. Next, we perform an in-depth quality assessment of HyDrop v2 generated data across different model organisms, i.e., mouse and fly, for commonly used bioinformatic tools and novel deep learning applications.

### HyDrop v2 captures the same cellular diversity in the mouse motor cortex as 10x v2

To evaluate the quality of the data generated with HyDrop v2, we generated scATAC-seq datasets on two complex tissues: the mouse cortex (Figure 1e) and the last four hours of *Drosophila melanogaster* embryo development (Figure 4a). To evaluate bead variability, we performed 35 experiments on frozen adult mouse cortex recovering a total of 110,547 cells after filtering (See Methods). For downstream analysis, the data integrates seamlessly with 10x (v1 and v2) and HyDrop v1 data (Figure S2a, c). With the combined data of 141,010 cells, we recover all major cell types of the mouse cortex (Figure 2a), annotated based on cell-type specific regions from two published data sets^12,24^.

**Fig. 2.**
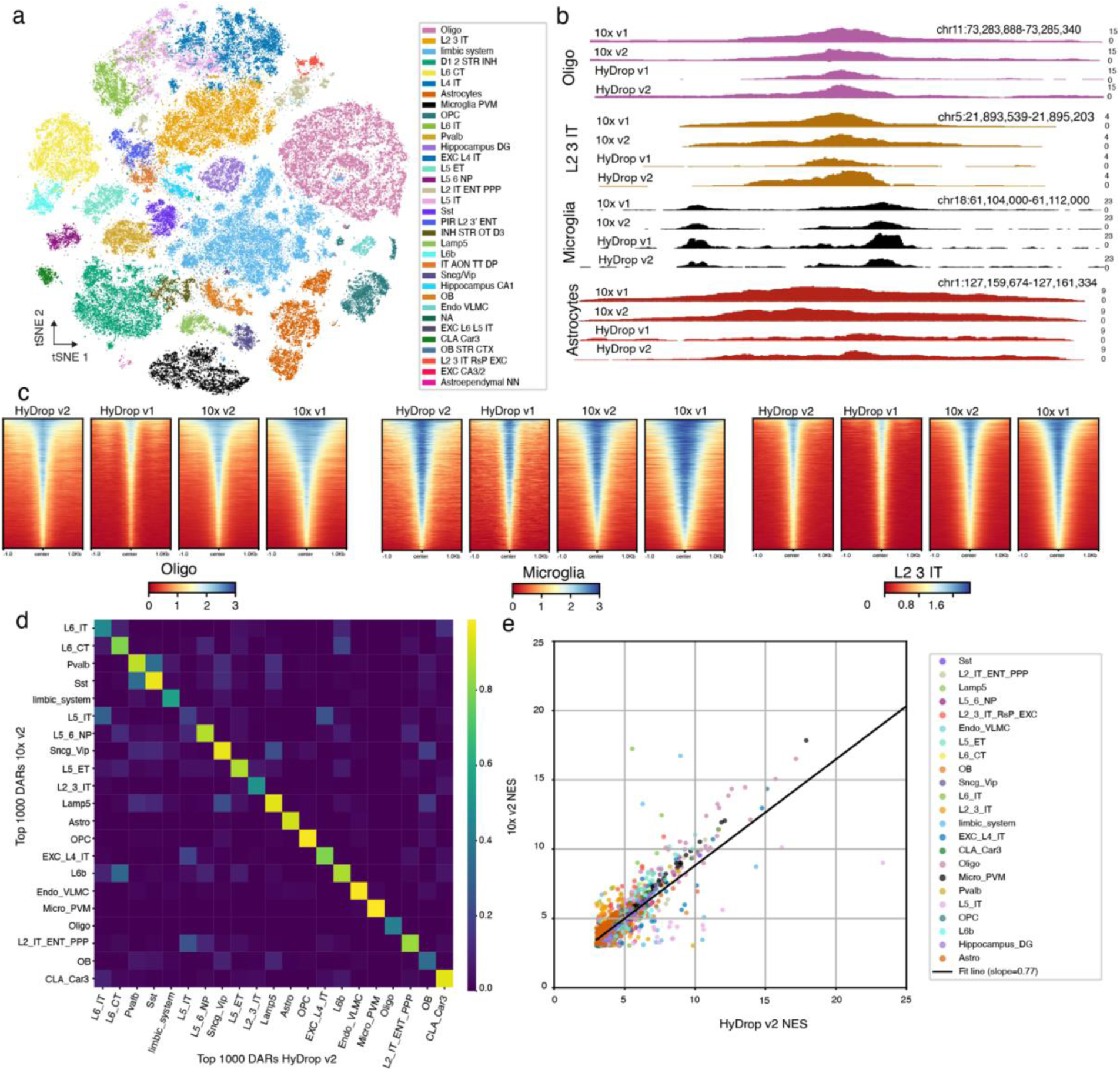
Differential accessible regions in HyDrop v2 mouse cortex replicate regions identified by 10x. **a,** *t*-distributed stochastic neighbor embedding (tSNE) of all 141,010 cells across 44 experiments (2 experiments 10x v1 with a total of 10,177 cells, 2 experiments 10x v2 with a total of 13,123 cells, 5 experiments HyDrop v1 with a total of 7,163 cells, 35 experiments HyDrop v2 with a total of 110,547 cells) colored by cell type, batch corrected for the used technique. Data points were randomly shuffled before plotting. **b**, Genome tracks of cell-type specific DARs down sampled to the lowest cell number present for each technique, 985 cells in oligodendrocytes, 469 cells in L2 3 IT neurons, 509 cells in microglia, and 998 cells in astrocytes. The shown genome tracks are normalized for the fragment count. **c**, carrot plot showing the regions of accessible chromatin ±1 kb around the center of DARs (logFC ≥ 1.5), sorted by the highest (blue) to lowest (red) accessibility for oligodendrocytes, microglia, and L2 3 IT neurons across the four different techniques. **d**, Heatmap of percental overlap of top 1000 DARs per cell type of 10x v2 and HyDrop v2 data. **e**, Scatterplot of normalized enrichment score of cell-type-specific transcription motifs, dots represent a common motif between HyDrop v2 and 10x v2 data colored per cell type as shown in a. DAR: differentially accessible region, NES: normalized enrichment score.

To compare cell-type features in-depth, we calculated differentially accessible regions (DARs) of each cell type per technique (HyDrop v2, HyDrop v1, 10x v2, 10x v1). HyDrop v2 reveals a similar fragment coverage as 10x v1 and v2 data in example cell-type specific DARs (Figure 2b) as well as across all cell-type specific DARs (Figure 2c). Only focusing on the 1,000 highest scoring DARs, i.e., regions yielding important signature features of each cell type^19^, reveals a high percental overlap between the regions recovered by HyDrop v2 and 10x v2 (Figure 2d): The mean overlap across cell types is 76.63% between 10x v2 and HyDrop v2 with a maximum of 98.8% recovering the same regions in microglia. Similarly, the 1,000 strongest DARs in HyDrop v1 show a mean overlap of 70.35% overlap with 10x v2 as well (Figure S2d).

Next, we evaluate the biological information covered in the cell-type specific DARs per technique using motif enrichment analysis (pyCisTarget^10^). The recovered motifs of 10x v2 and HyDrop v2 correlate highly on their normalized enrichment score (NES, r=.77) per cell type (Figure 2e), while HyDrop v1 performed slightly lower (NES r=.74, Figure S2e). A close-up of enrichment scores of exemplified cell types can be found in Figure S2f.

### Sequence-to-function model identifies cell type-specific enhancers in the adult mouse cortex

Sequence-based deep learning models trained on scATAC data have been shown to identify cell-type specific sequence features in the genome^12,13^. Published models thus far have been trained on data generated by commercial platforms such as 10x genomics^12–14^. Here, we investigated whether scATAC-seq data generated by HyDrop v2 can be used to train performant S2F models. To this end, we first clustered all HyDrop v2 mouse cortex data, integrating seamlessly without batch correction (Figure 3a, 1h). Next, we trained S2F models using the CREsted package^22^ on HyDrop v2 data predicting peak heights across cell types. To evaluate the model performance, we compared the HyDrop v2-based S2F model to a published model trained with 10x multi-ome data and which outperformed other S2F models earlier (see BICCN challenge^12^ for details). When comparing the correlation of the prediction versus ground truth peak heights, both models closely predict the peak height as seen in their respective data sets (Figure S4a, b). The HyDrop v2 model outperforms the 10x model in the class-wise comparison on the full test set, while the 10x model scores slightly higher in the cell type-specific test set (Figure 3b) as classically seen for fine-tuned models^12,21^. In the region-wise comparison, HyDrop v2 closely follows the 10x model.

**Fig. 3.**
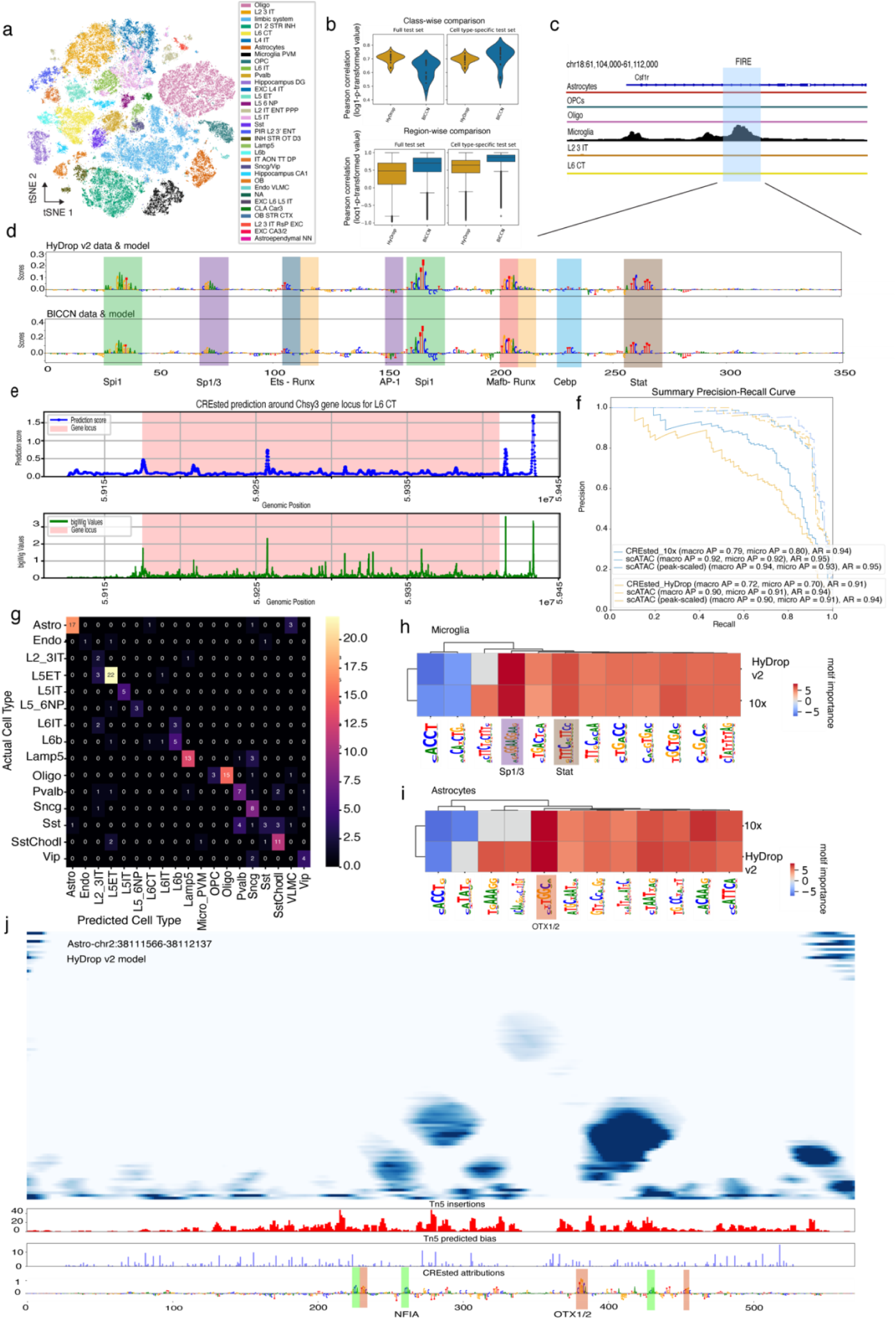
Hydrop v2 produces comparable training data for cutting-edge deep learning tools as public 10x data set. **a**, *t*-distributed stochastic neighbor embedding (tSNE) of all 110,547 cells generated with HyDrop v2 (35 experiments) colored by cell type, no batch correction. Data points were randomly shuffled before plotting. **b,** Model comparison of sequence models trained on HyDrop v2 data (orange) and the BICCN (10x) public data set (i.e., 10x multi-ome data in blue). The accuracy of the predictions evaluated based on the ground truth of region accessibility is compared on class-wise (top) and region-wise level (bottom) for both the full test set in the data and cell type-specific test sets. **c**, Genome tracks of cell-type specific chromatin accessibility of the FIRE enhancer (mm10 chr18:61108475-61108975), regulating Csf1r expression in microglia. **d**, Nucleotide contribution score of the FIRE enhancer from a sequence model trained on HyDrop v2 data (top) and a model trained on 10x data. Previously described TF binding sites are highlighted for the corresponding TF. **e,** Prediction by the HyDrop v2 sequence model on HyDrop v2 data of chromatin accessibility for L6 CT (top) around the Chsy3 gene locus compared to actual chromatin accessibility plotted from the corresponding bigwig file (bottom). **f**, Specificity prediction for *in vivo* enhancer activity of validated mouse cortex enhancers by Ben-Simon et al. (2024) shown for the HyDrop v2 and BICCN (10x-based) model. **g,** Heatmap of enhancer indicated by Ben-Simon et al. (2024) identified as cell-type specific enhancer of the corresponding cell type by the HyDrop v2 model. **h**, **i,** Evaluation of motif importance scores of highly relevant regions for cell type identity indicated by the BICCN (10x) data-based model and HyDrop v2 data-based model for microglia (**h**) and astrocytes (**i**) with motifs shown in d and j, respectively, are identified and highlighted. **j,** Multiscale footprint of the region (mm10 chr2:38111566-38112137) of HyDrop v2 data only. Bottom tracks show Tn5 insertion, the predicted Tn5 bias and the nucleotide contribution scores based on the HyDrop v2 sequence model. Previously described TF binding sites are highlighted for the corresponding TF.

To investigate whether the HyDrop v2 model captures relevant biological information, we evaluate its peak height predictions and sequence explanations: Similar to previously published models^13^, the HyDrop v2-based model recovers the correct microglia TFBS in the FIRE enhancer that replicate the findings of the 10x model (Figure 3c-d). Furthermore, the HyDrop v2 model can correctly predict chromatin accessibility in an unseen locus such as the *Chsy3* gene (Pearson correlation coefficient: 0.59), a marker gene of L6CT neurons (Figure 3e top). To investigate if the HyDrop v2 model can identify enhancers across major mouse cortex cell types, we compared its performance to a suite of enhancers earlier evaluated *in viv*o^25^. The HyDrop v2 model assigns the validated enhancer sequences with an average precision (AP) of 0.70 to the correct cell types (Figure 3f), while the optimized 10x model achieves an AP of 0.80. The prediction of enhancer activity based on scATAC information is comparable between HyDrop v2 and 10x data (Figure 3f, AP: 0.91-0.93). A similar observation is made when looking at the separate enhancers per cell type (Figure S3).

To further investigate the sequence explainability of each model for cell type-specific enhancers, we performed de novo motif discovery using *tfmodisco lite*^26^ through the CREsted package^22^. Per trained model (10x vs HyDrop v2), we identified enriched patterns, alongside their individual instances (called seqlets). For the highest and lowest scoring patterns with a contribution score cut-off 4.25, we find the same patterns in cell types in 10x and HyDrop v2 data for astrocytes and microglia (Figure 3h, i) just as for several neuronal cell types (Figure S4g-j).

S2F models trained on commercial scATAC data have been shown to accurately identify cell-type specific regions and predict TF binding sites^13^. Here, we show that a model trained on non-commercial data generated with our open-source HyDrop v2 protocol achieves similar performance to an optimized and fine-tuned 10x model and, importantly, can recover cell type-specific regions and TF-binding sites. To investigate whether the slight difference in performance can be attributed to model variability or a difference in training data quality, we trained two models with 10-fold cross-validation on 10x v2 and HyDrop v2 of the *Drosophila* embryo generated with the same library preparation protocol, data processing, and model architecture (see below).

### Cellular diversity in Drosophila embryos is captured comparably with HyDrop v2 and 10x v2

Apart from the mouse cortex samples, we compared the optimized HyDrop v2 protocol on another complex biological system, the developing fruit fly. *Drosophila melanogast*er is widely used for developmental research^27^ but is rarely included in larger benchmarking studies of scATAC techniques^19,20^. Here, we compare in-house generated HyDrop v2 and 10x v2-based data of the last four hours of *Drosophila* embryo development (stage 17)^27^. Additionally, we compare the quality of these two microfluidic platforms to public data from the *Drosophila* embryo of the same age^27^ generated with sciATAC-seq3, a plate-based assay using combinatorial indexing^28^ (Figure 4a). We generated a new embryo atlas with 10x v2 and HyDrop v2, with a total of 607,340 cells (HyDrop v2: 340,604 cells, 10x v2: 266,736 cells, Figure 4b). We annotated this new atlas based on the region accessibility of DARs of the public embryo atlas^27^ combined with annotated activity patterns of *in vivo* tested enhancers^17^. HyDrop v2 reveals a similar fragment coverage as 10x v2 data in, cell-type specific DARs (Figure 4d top, S5d) as well as across all cell-type specific DARs (Figure 4e left, S5e). sci-ATAC data exhibits a wider range of accessibility across regions compared to 10x v2 and HyDrop v2 data (Figure 4d, e and S5d,e). However, when focusing on the cut sites of Tn5 (Figure 4d bottom, S5d), sciATAC resembles the Tn5 insertion profile observed in 10x v2 and HyDrop v2 data, yielding a more distinct identification of accessible region across all cell-type specific DARs (Figure 43 right, S5e). Possibly due to the larger fragment size covering wider regions of the genome (Figure 4d, S5c-d), sciATAC has a significantly higher fraction of reads in peaks (FRIP) than HyDrop v2 and 10x v2 (Figure 4c left, H(2) = 37,728, p<.001) while both 10x v2 and HyDrop v2 score higher on TSS enrichment (Figure 4c middle, H(2) = 172,030, p<.001) and the number of unique fragments per 12k reads per cell (Figure 4c left, H(2) = 48,846, p<.001). The in-house generated data was downsampled to 12k reads per cell to not overestimate the read compared to sciATAC^27^. Similar to mouse HyDrop v2 data, the slightly higher ability to recover unique fragments of 10x v2 versus HyDrop v2 is reflected by its more efficient sequencing at the same depth (Figure S5g). When investigating the depth of biological information, we found that the most important 1,000 DARs identified by HyDrop v2 or 10x v2 overlap with a mean of 80.57% across cell types (Figure 4f), just as in the mouse cortex data. When performing motif enrichment analysis^10^, we observed a strong correlation (r=.95) of the enrichment score of individual enriched motifs between 10x v2 and HyDrop v2 data (Figure 4g) and, to a slightly lesser extent, in 10x v2 and sciATAC (Figure S5h, r=.92).

**Fig. 4.**
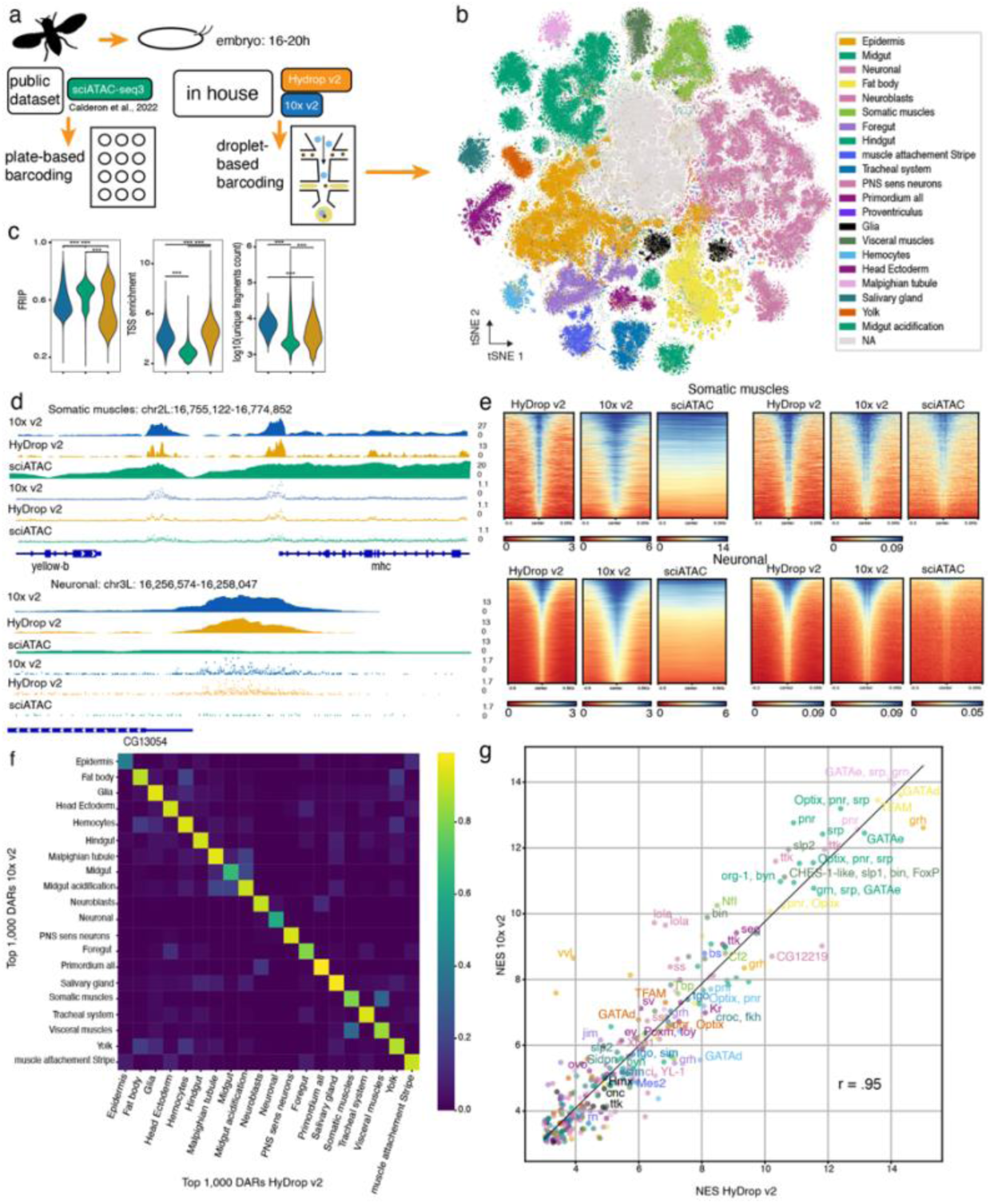
HyDrop v2 captures cellular diversity in developing Drosophila melanogaster. **a,** Overview of study design. Drosophila melanogaster embryos were collected 16-20h after egg laying. In-house experiments were performed with 10x v2 chemistry on the 10x Chromium device, while HyDrop v2 beads were used for the open-source data generation. sciATAC-seq3 data (fragment files for the plate-based essay) were retrieved from Calderon et al., (2022). **b**, *t*-distributed stochastic neighbor embedding (tSNE) of all 607,340 cells generated with HyDrop v2 (340,604, 18 experiments) and 10x v2 (266,736 cells, 5 experiments) colored by cell type, batch correction according to wet lab protocol used to generate data. Data points were randomly shuffled before plotting. **c,** Violin plots of quality measurements (TSS enrichment, FRIP, log fragment counts) compared between 10x v2, sciATAC-seq3, and HyDrop v2 data**. d,** Genome tracks (upper) and cut site insertion of Tn5 (lower) of cell-type specific DARs down sampled to the smallest count present for each technique, 1,861 cells in neuronal cells and 8,916 cells in somatic muscle cells. The shown genome tracks are normalized for the fragment count. **e**, Carrot plot showing the regions of accessible chromatin ±0.5 kb around the center of DARs sorted by the highest (blue) to lowest (red) accessibility for neuronal cells and somatic muscle cells in 10x v2, HyDrop v2, and sciATAC data. DARs are calculated on fragment coverage (left) and Tn5 cut sites (right) for each technique. **f**, Heatmap of percental overlap of top 1,000 DARs per cell type of 10x v2 and HyDrop v2 data. **g,** Scatterplot of normalized enrichment score of cell-type-specific transcription motifs, dots represent a common motif between HyDrop v2 and 10x v2 data colored per cell type as shown in a, the labels are shown for the top three motifs per cell type. DAR: differentially accessible region, TSS: Transcription start side, FRIP: fraction of reads in peaks.

Just as for mouse cortex scATAC, we show that HyDrop v2 offers a comparable alternative to 10x v2-based data in the developing fruit fly. Overall, HyDrop v2 *Drosophila* embryo data quality has a slightly lower sensitivity *per cell* compared to 10x v2 but due to the low cost, larger atlases can be constructed and at the atlas level, the quality of detecting DARs and enriched motifs become equivalent.

### Essential regions in validated Drosophila enhancer recovered by Hydrop-v2 based deep learning model

Apart from investigating data quality in terms of the common bioinformatics analyses, e.g., DARs, and motif enrichment analysis, we trained separate S2F sequence models on the *Drosophila* embryo data for 10x v2 and HyDrop v2 (Figure 5a). Again, the fly models were trained using the CREsted package, although for fly we applied additional filtering on the standardized peak heights (see Methods). Both, the model based on 10x v2 and the model based on HyDrop v2 score identical on region and class-wise prediction of databased accessibility (Figure S6a-c).

**Fig. 5.**
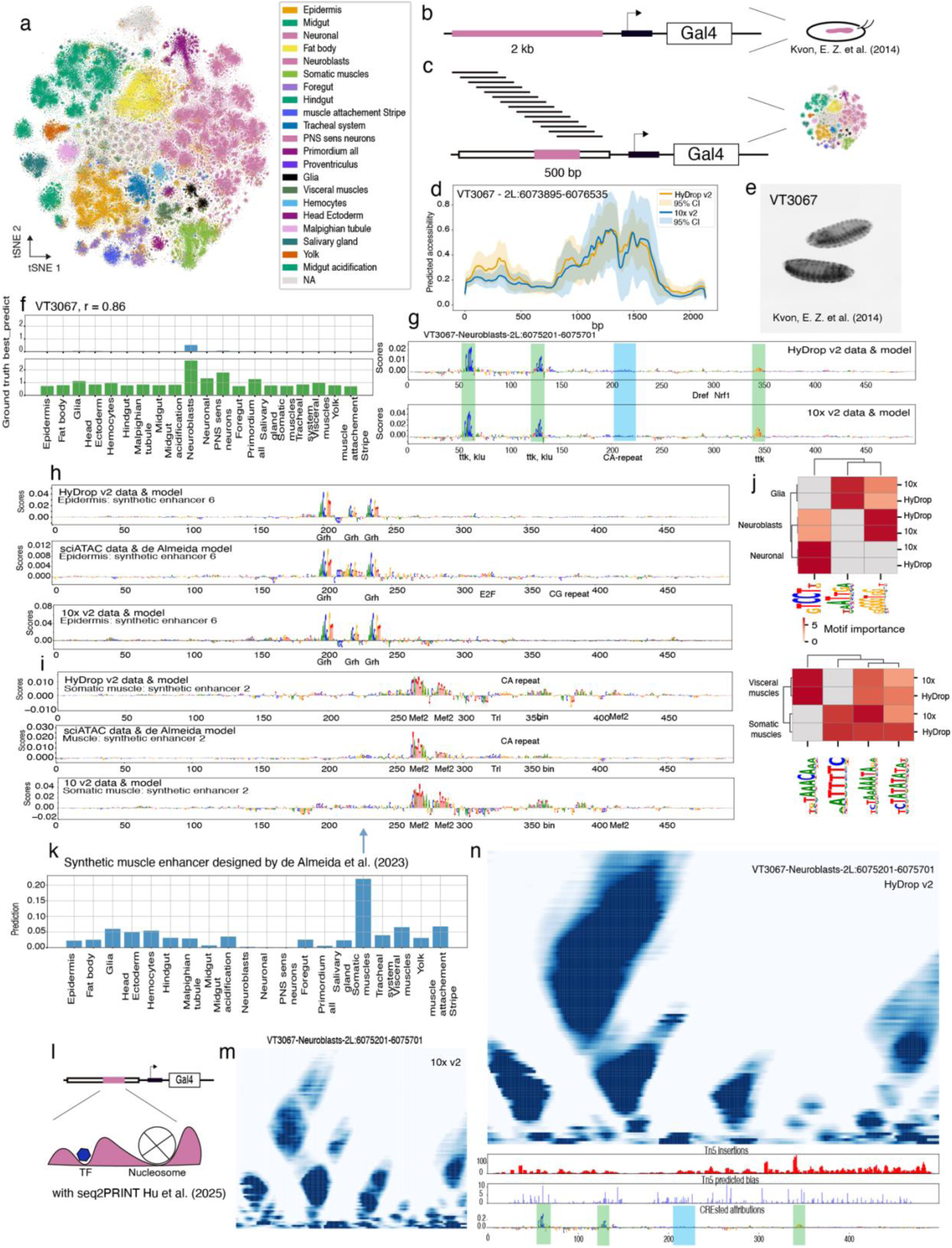
HyDrop v2 data enables training of sequence-based Deep Learning model to uncover cell-type specific transcription factor motifs and footprints in Drosophila embryo. **a**, *t*-distributed stochastic neighbor embedding (tSNE) of 340,604 cells (18 experiments) HyDrop v2 colored by cell type, no batch correction is applied. Data points were randomly shuffled before plotting. **b,** VDRC library of transgenic flies with ∼2kb enhancers (pink) upstream of minimal promotor and Gal4 reporter. 2,693 enhancers stage 15-16 evaluated *in vivo* by Kvon et al. (2014). **c,** Scanning of ∼2kb enhancers with sliding window (10bp shift) of 500 bp evaluated on cell types seen in a. **d,** Predicted accessibility of 500 bp windows of VT3067 enhancer from VDRC library. Both, the predictions based on the HyDrop v2 and 10x v2 data are shown with confidence intervals of 10-fold replicate models. e, *In situ* hybridization of VT3067-Gal4 reporter embryos stage 15 with antisense Gal4 probe by Kvon et al. (2014). **f,** Prediction (top) versus ground truth (bottom) of 500bp region within 2kb long VDRC enhancer VT3067 active in neuroblasts. **g,** Nucleotide contribution score of the 500 bp within VT3067 enhancers (VDRC fly line) trained on HyDrop v2 data (top) and 10x v2 data (bottom). Previously described TF binding sites are highlighted for the corresponding TF. **h,** Nucleotide contribution score of the 500 bp of synthetic enhancer designed to be active in epithelial cells based on sciATAC data by de Almeida et al. (2023): HyDrop v2 and 10x v2 model predictions shown for epithelial cells versus predictions by the original model for somatic muscle cells data. Previously described TF binding sites are marked for the corresponding TF. **i,** Nucleotide contribution score of the 500 bp of synthetic enhancer designed to be active in muscle cells based on sciATAC data by de Almeida et al. (2023): HyDrop v2 model predictions shown for somatic muscle cells (top) versus predictions by the original model for general muscle cells data (bottom). Previously described TF binding sites are highlighted for the corresponding TF. **j,** Evaluation of motif importance scores of highly relevant regions for cell type identity indicated by the 10x v2 data-based model and HyDrop v2 data-based model for neurons, glia, neuroblasts, and visceral muscles compared to somatic muscles. An importance threshold of 5 was handled. **k,** Prediction of tissue accessibility of synthetic enhancer designed for muscle cells in 10-12h embryos. Enhancers were designed by de Almeida et al. (2023) and scored with the HyDrop v2 model. Accessibility is specifically predicted for somatic muscle cells in the HyDrop v2 data. **l,** Schematic overview of possible footprinting in selected 500 bp region of VDRC library enhancers. Footprinting is performed with seq2PRINT suit by Hu et al. (2025). **m,** Multiscale footprint of 500bp region in VT3067 in 10x v2 data. **n,** Multiscale footprint of 500bp region in VT3067 in HyDrop v2 data only. The bottom track shows Tn5 insertion, the predicted Tn5 bias, and the nucleotide contribution scores based on the HyDrop v2 sequence model. Previously described TF binding sites are highlighted for the corresponding TF. VDRC: Vienna Drosophila Resource Center

To evaluate the ability of our S2F models to capture biologically relevant information, we made use of validated enhancers^17^ analogous to the evaluation of the mouse models. Kvon et al. (2014) characterized 3,557 developmental enhancers *in vivo* according to tissue specificity of enhancer activity. This valuable enhancer database contains 2,693 enhancers assayed in stages 15-16 of which 80 are highly active with a specificity of maximally five tissues^17^. As the characterized enhancers have a length of approximately 2kb (Figure 5b), we scanned the regions with a sliding window of 500 bp (Figure 5c) in a 10 bp shift to determine the region that is the most relevant for the enhancer activity in the respective tissues in (Table S1) according to our S2F models. Both HyDrop v2 and 10x v2, as well as their respective 10-fold cross-validation models (shown as confidence intervals), identify the same 500 bp regions as primarily accessible across 2kb enhancers (Figure 5d, S6d). Enhancer VT3067, mainly active in the ventral nerve cord (VNC, Figure 5e), is predicted to be accessible in neuroblasts (NBs, Figure 5f), linking activity to enhancer accessibility of the 500 bp window with the highest predictive value. Also, in other tissues, the observed *in vivo* activity^17^ corresponds to the predicted accessibility in the matching cell type based on the highest scoring 500 bp window of the respective enhancer using the HyDrop v2 model (Figure S6f,h for midgut cells, S6j,l for foregut cells, S6n,p for epidermis).

In the developing *Drosophila*, NBs type 1 and 2 are found in the VNC giving rise to neurons and glia^29^ with the first wave of neurogenesis at the end of embryo development^30^. The nucleotide contribution scores of both HyDrop v2 and 10x v2, indeed predict motifs of NB-specific TFs such as *klumpfuß* (*klu*)^31^ and *tramktrack* (*ttk*)^29^ (Figure 5g). Interestingly, performing *tfmodisco lite* identifies the *ttk* motif (TCCT, ACCC) in NB and neurons but not glia. This is in line with well-established findings of *ttk* repressing glial fate in differentiating NBs^29^. Notably, the observed TF-motifs such as *ttk* or aspecific CA repeats match the reported motifs by the earlier central nervous system (CNS) S2F model trained on a combination of sciATAC data and enhancer activity data ^15^, as NBs occur in the CNS of the developing *Drosophila* embryo^29^. Comparing somatic and visceral muscle cells, we observe different motifs with, e.g., *dorsal* only being predicted in somatic muscle cells (Figure 5j). The ability to identify different motifs in closely related tissues underlines the power of HyDrop v2 S2F to models to capture biological relevance.

The authors of the previously published *Drosophila* embryo S2F models designed synthetic enhancers for five tissues (CNS, brain, muscle, epidermis, and gut) using one model per tissue as a guiding oracle^15^. Per tissue, the activity of eight enhances was validated *in vivo*^15^. To compare our HyDrop v2 and 10x v2-based models with the sci-ATAC-based models, we evaluated the synthetic sequences designed by de Almeida et al. (2023). Due to the higher resolution in the annotation of our data and a model design that includes all cell types, we could specify that synthetic enhancer 1^15^, which is designed for muscle cells, is indeed specific for somatic muscles across the *Drosophila* embryo (Figure 5k). Additionally, the HyDrop v2 model recovers all motifs indicated by the sciATAC model, while the 10x model is missing one *Trithorax-like* (*trl*) motif (Figure 5i). Similarly, nucleotide contribution scores of synthetic epidermis enhancer 6 (Figure 5h, S7c) show the typical *grainyhead* (*grh*) motif; a TF essential to epithelial cell fate^32^. The multi-class set of our HyDrop v2 S2F model allows for scoring synthetic enhancer activity across tissues. Synthetic enhancer 8, originally designed for epithelial cells, is mainly predicted to be accessible in glial cells based on our model (Figure S7d). Evaluating the nucleotide contribution indicates *repo* motif instances, being indicative for glia^29^, in both 10x v2 and HyDrop v2 data. The mix of motifs from different tissues in this enhancer identified by our models might explain the lower predicted activity in the original sciATAC-based model^15^.

Notably, not all of the previously designed synthetic enhancers showed (matching) *in vivo* activity^15^. Especially the brain and gut enhancer design is mentioned to be cumbersome. While the brain is a highly heterogeneous tissue itself, the gut poses the difficulty of being developed from the ectoderm (foregut and hindgut) and endoderm (midgut)^33^, making a targeted synthetic enhancer design possibly less straightforward compared to tissues with one developmental origin. By including a large variety of tissues into one single model, we find that the synthetic enhancers that are not specifically active in the target tissue^15^ (generally speaking, the gut) are also predicted to be active in other tissues outside the gut (Figure S73). This offers a possible explanation for the lack of specificity observed by de Almeida et al. (2023).

Overall, we show that HyDrop v2 produces data of high quality that can be used to train multi-class S2F models, revealing biologically relevant features that closely resemble findings from 10x S2F models. Possible variations between models can be attributed to model variability, as shown in the 10-fold cross-validation (Figure 5d). Additionally, in *Drosophila*, we identified the core 500 bp of *in vivo* validated enhancers and replicated the findings of synthetic enhancer design in the *Drosophila* embryo.

### HyDrop v2 matches transcription factor footprints found in 10x data in validated mouse and *Drosophila* enhancers

Next, we benchmarked the performance of HyDrop v2 data against 10x data to predict TF footprints. To this end, we applied the recently developed seq2PRINT framework^34^. seq2PRINT uncovers footprints of DNA-protein interaction in scATAC and bulk ATAC data after correcting Tn5 bias with a convolutional neural network^34^. The use of a species-specific Tn5 bias model permits the evaluation of possible footprints of nucleosomes and smaller proteins, such as transcription factors, in *Drosophila* and mouse cortex data. We first investigated whether the Tn5 bias differs between HyDrop v2 and 10x data, by comparing the Tn5 cut sites across techniques, species, and cell types. Tn5 cut sites obtained by HyDrop and 10x per cell type appear uniformly flanking the center of the peak in the accessible regions in mouse (Figure S4k-m) and fly (Figure 4d, S5e). The footprint around the center of the peak reflects the seq2PRINT-given Tn5 bias consistently in each cell type (Figure S4k-m, S5f). Interestingly, the footprint (the double vertical line) tends to be more visible in *Drosophila* due to the higher Tn5 concentration - relative to the small genome of 180 Mb^35^ - resulting in smaller fragments. Consequently, performing the library preparation with the same Tn5 concentration as in mouse (see Methods) results in a higher insertion ratio of Tn5, leading to shorter fragments in *Drosophila* (Figure S5c) compared to mouse (Figure S1g). Importantly, just as in mouse, the Tn5 bias does not differ between HyDrop v2 and 10x v2 in *Drosophila* samples.

After correcting for possible Tn5 bias, we apply seq2PRINT in mouse (HyDrop v2 data and 10x-multi-ome data^12^ from the 10x mouse model) and *Drosophila* (HyDrop v2 data and in-house 10x v2 data). In mouse, seq2PRINT identifies possible TF footprints that align with the predicted motifs based on the S2F model in the *in vivo* validated enhancers in astrocytes (Figure 3j, S4e), L5 ET neurons (Figure S4c,d), and oligodendrocytes (Figure 4f). Larger footprints indicating nucleosome binding coincide with the depletion of motifs. Importantly, the predicted footprints in HyDrop v2 data largely correspond to 10x data-observed footprints.

In the *Drosophila* embryo, we focused on the 500 bp window with the highest prediction score of the *in vivo* validated enhancers. We observed possible footprints coinciding with motif locations predicted by the S2F models that match between HyDrop v2 and 10x v2-based seq2PRINT predictions for NBs (Figure 5m,n), midgut cells (Figure S6e,g), foregut cells (Figure S6i,k), and epidermis cells (Figure S6m,o).

In conclusion, the improvements from HyDrop v1 to v2 chemistry result in reliable data generation of large-scale scATAC data sets in complex tissue covering various cell types. Our results show that HyDrop v2 data enables building large-scale atlases with similar resolution to the gold standard commercial microfluidic platform in mouse and fly achieving comparable findings in commonly used bioinformatic applications and novel deep learning tools.

## Discussion

Identifying cell-type-specific regulatory sequences from scATAC-seq data has become a driving force behind the unraveling of gene regulatory networks but the cost-effective generation of comprehensive scATAC-seq atlases is often a bottleneck. While S2F models offer powerful ways to identify CREs such as enhancers, their performance strongly depends on the quality and modality of training data^12^. Currently, scATAC-seq data acts as the gold standard for S2F model training^12,13,21,34^. To address the need for accessible, high-quality, and cost-efficient scATAC-seq data generation, we improved upon our previously established open-source microfluidic platform HyDrop (now HyDrop v2), enabling consistent generation of large-scale data for improved DL-based enhancer identification.

First, we generated large atlases of the mouse motor cortex with 110k high-quality cells and the developing embryo of age 16-20h after egg laying of *Drosophila melanogaster* with 340k cells using the novel HyDrop v2 platform. The costs of generating the library of, for example, the mouse cortex across 35 experimental “runs” accumulate to merely 1,080.60 euros compared to an equivalent of 10x v2-based library preparation for 15,330.70 euros, excluding the sequencing cost. With the improved workflow, a single trained technician can perform 16 HyDrop v2 runs in 1.5 days. Thus, generating >100k cells of mouse cortex library becomes feasible within a few days, making large-scale data generation with HyDrop v2 possible at low cost and scale.

After library preparation, the batches are sequenced to a desired read depth per cell. Here, HyDrop v2 libraries reach a comparable number of unique fragments per cell to 10x data but require a higher sequencing depth (i.e., having a lower sequencing efficiency). Nevertheless, we found that the differentially accessible regions per cell type captured in the HyDrop v2 atlases are equivalent to the 10x-recovered regions in mouse and fly, even after equilibrating for sequencing depth (i.e., raw number of reads per cell). This confirms earlier findings showing that the number of DARs does not depend on sequencing depth^19^. HyDrop v2 libraries already deliver comparable quality to 10x data at a sequencing depth of 36kRPC for mouse and 12kRPC for the *Drosophila* embryo. HyDrop v2 data integrates seamlessly with data from other microfluidics-based platforms, and the assay can therefore also be used for protocol optimization before investing in a major experiment. Eventually, data generated with different droplet-based platforms can be combined for large-scale atlases across species.

Next, we evaluated the performance of HyDrop v2-based atlases in two deep learning tools: ready-to-use S2F modeling^22^ and seq2PRINT^34^. As the mouse cortex is well represented in available S2F models, we chose a published 10x-based model^12^ to compare our default HyDrop v2 model. Across the major mouse cortex cell types, the HyDrop v2 model learned relevant sequence features and identified candidate motifs comparable to the 10x-based model. In predicting the activity of *in vivo* validated enhancers^35^, HyDrop v2 model uncovers most tested enhancers while we see a slightly higher performance of the 10x model which was, however, optimized in model architecture and finetuned.

Previous benchmarking studies have shown that method comparison is not trivial when many factors (e.g., library preparation, barcoding platform, sequencing depth, processing, annotation resolution) have to be taken into account^19^. Here, the variability of models with and without finetuning adds an additional variable. Besides, the difference observed in the performance of HyDrop v2 and 10x S2F models may be due to differences between the data sets, for example, the 10x dataset had been curated to include only motor cortex cell types, while the HyDrop dataset does not differentiate between endothelial cells or vascular leptomeningeal cells (VLMC), vasoactive intestinal polypeptide reactive interneurons (VIP) or their *SNCG*-positive subtype, and Sst (somatostatin-expressing) cells and their *Calb2*-positive subtype (SstChodl). These cell types were investigated separately by the test enhancer set; thus, the difference in annotation resolution might underlie the slightly lower of the HyDrop v2 model. That makes it even more remarkable that the HyDrop v2 S2F model with a default CREsted architecture scores almost comparable to the 10x model that performed among the best across other earlier tested models in a worldwide community challenge^12^ emphasizing the usability of HyDrop v2 to train deep learning models.

To, nevertheless, address the sources of variability, we generated parallel datasets in terms of library preparation protocol, data processing, and annotation of the *Drosophila* embryo. We then trained analogous models with 10-fold cross-validation to also factor in the model variability of S2F models. Additionally, most deep learning studies^12,13,21^ focus on cell lines or tissues of larger model organisms such as mice, resulting in a lack of evaluation of tools applied to data from non-vertebrate model organisms. Here, we trained S2F model on HyDrop v2 and 10x v2 data of the *Drosophila* embryo with (1) data generated with the same library preparation protocol, (2) the same model architecture and (3) comparable cell numbers. With the 10x v2 and HyDrop v2 models, we were equally able to pinpoint the enhancers within *in vivo* tested regions of ∼2kb size, in late-stage embryos. The 10-fold cross-validation shows that there is indeed variability between S2F models in cell type-specific accessibility prediction but we, nevertheless, observe a large agreement between the cross-validations in enhancer region selection. With motif enrichment analysis and deep learning-based *de novo* motif discovery, we replicate earlier findings, underscoring the ability of HyDrop v2 data to uncover biological relevant information comparable to 10x just as in mouse.

Additionally, we compared both our *Drosophila* S2F models (HyDrop v2 and 10x v2) to a previously published model that was trained on sciATAC-seq3 data, a plate-based assay^15,27^. The five models by de Almeida et al. (2023), one per target tissue, had been finetuned with additional *in vivo* enhancer assays, as opposed to our models that are trained on scATAC-seq data only. In mouse, it has been shown that the recent advances in S2F model design allow for improved models without the need for additional finetuning based on additional data modalities^12^. Models trained on additional data modalities in mouse are evaluated elsewhere^12^. Besides differences in model design, de Almeida et al. (2023) are trained on data of 10-12h of embryo development^15^, while our models are trained on a slightly later developmental stage of *Drosophila* development. Additionally, the sciATAC-based models focus on CNS, brain only, epithelial cells, gut cells, and muscle cells while our models include the entire embryo. As especially CNS development undergoes major changes (i.e., the embryonic wave of neurogenesis) in the last hours of embryo development^30^, we do not include the CNS and brain models by de Almeida et al. (2023) in our comparison. The synthetic enhancer sequences designed for gut, epidermis, and muscle tissue^15^ are identified as target sequences matching the respective cell types in our multi-class model. For muscle, the annotation in our data allows for specifying the synthetic enhancer for the muscle to be active in the somatic muscle tissue, which is also confirmed by the enhancer’s *in vivo* validation score^15^. Here, also *de novo* motif discovery driven by our S2F HyDrop v2 model uncovers different motifs important in somatic versus visceral muscles.

It becomes apparent that the larger fragments of the sciATAC data cover a wider range of regions compared to 10x v2 and HyDrop v2 when using the same pre-processing. Regarding DARs and motif enrichment, sciATAC only slightly underperforms or achieves equal results as 10x and HyDrop v2. The larger fragment size of sciATAC, however, might underlie the need for finetuning of the sciATAC-based S2F model by de Almeida et al. (2023) and the difficulty of integrating sciATAC data with HyDrop v2 and 10x v2 data of the same developmental time stage in our hands. In contrast to sciATAC, HyDrop v2 and 10x v2-based data in *Drosophila* resemble the mouse genome coverage used for successful model generation in mouse^12,13^, cell lines^36^, or the adult fly^36,37^. This finding underscores the need for a priori evaluation of the input data for deep learning models.

Besides investigating the performance of HyDrop v2 data in bioinformatics tools such as motif enrichment and more advanced S2F models, we compared footprints found in 10x and HyDrop v2 data for mouse and fly. In both model organisms, seq2PRINT^34^ identifies similar multi-scale footprints, highlighting the wide range of applications that can be reliably performed with data generated with the improved HyDrop v2 assay.

Additional comparisons of other scATAC platforms, such as microwell encapsulation of single cells (BD Rhapsody) or well-based combinatorial indexing strategy (Scale Bioscience) for deep learning purposes might be needed to benchmark S2F modeling across a diverse range of experimental techniques. Another plate-based platform called SPATAC is used for zebrafish^38^ data generation. In the recent CREsted implementation, the authors show that SPATAC data yields enough information to train S2F models and synthetic enhancer design^22^. To our knowledge, no data for *Drosophila* or mouse cortex generated with SPATAC is available to compare to droplet- and other plate-based platforms.

With HyDrop v2, we offer an accessible, scalable, and low-cost method to generate large-scale scATAC-seq datasets suitable for training read-to-use deep learning tools (S2F models and footprinting) without the immediate need for further finetuning. We demonstrate that HyDrop v2 data reliably captures biologically relevant regulatory features and can achieve predictive performance comparable to commercial tools such as 10x Genomics. In combination with the easy-to-interpret bioinformatic tools, HyDrop v2 makes gaining biological insights by generating high-quality and low-cost data scATAC more accessible to a wider range of researchers. The integrability of HyDrop v2 data with 10x Genomics data allows for cost-effective upscaling of data generation across species.

By providing an evaluation of droplet-based scATAC-seq strategies to generate high-quality training data, we contribute to the improvement of S2F models and other bioinformatics applications. We further exemplify the importance of close interactions between data generation technologies and machine learning in biology. The advances in S2F models to learn the enhancer grammar underlying cell type-specific CREs enable researchers to design synthetic enhancers^15,22,36^ targeting specific cell types only. Improving the data generation of large-scale atlases will elevate the design of reliable synthetic enhancers that could be used to develop novel disease treatment strategies. With the increased availability of deep learning tools, future benchmarking studies may be needed to reevaluate newly available bioinformatics tools on their performance on large-scale datasets, ideally composed of data generated on different platforms for cost-effective use of resources. Our results underscore the importance of evaluating scATAC data across platforms, paving the way for future benchmarking studies to ensure broad generalizability and reproducibility of deep learning approaches in regulatory genomics.

## Data availability

The scATAC coverage bigwigs can be downloaded at https://ucsctracks.aertslab.org/papers/hydrop_v2_paper/. The sequencing data and count matrix are available at Gene Expression Omnibus accession number GSE293575. For mouse cortex data, this includes the public data for mouse cortex downloaded from 10x Genomics (https://www.10xgenomics.com/datasets/fresh-cortex-from-adult-mouse-brain-p-50-1-standard-1-1-0, https://www.10xgenomics.com/datasets/8k-adult-mouse-cortex-cells-atac-v1-1-chromium-x-1-1-standard, https://www.10xgenomics.com/datasets/8k-adult-mouse-cortex-cells-atac-v2-chromium-controller-2-standard) and earlier published data of De Rop et al. (2022) with raw data available at GSE175684. The sciATAC-seq data of the Drosophila embryo age 16-20h after egg laying were downloaded from Calderon et al. (2022). All processed fragment files are available at https://resources.aertslab.org/papers/hydrop_v2/. The S2F models can be found in the CREsted repository https://crested.readthedocs.io/en/stable/models/Hydrop/index.html.

## Code availability

A detailed explanation of the PUMATAC pipeline to process HyDrop v2 and 10x data can be found at https://github.com/aertslab/PUMATAC. All data analysis and model training scripts will become available at https://github.com/aertslab/. Detailed instructions on CREsted can be found at https://github.com/aertslab/CREsted and https://crested.readthedocs.io/en/latest/changelog.html. For Seq2PRINT (scPRINTER in python implementation), we refer to the tutorial found here https://github.com/buenrostrolab/scPrinter by Hu et al. (2025).

## Funding

This research was funded in part by Aligning Science Across Parkinson’s (ASAP-000430 and 1512 ASAP-025179) through the Michael J. Fox Foundation for Parkinson’s Research (MJFF); ERC AdG (101054387); CZI (DI2-0000000068); SBO (S005024N); FWO (G094121N, G044124N); VIB Tech Watch funding to S.P & S.A., FWO fellowship to H.D. (1168625N), FWO fellowship to F.D.R. (1S80920N), FWO PhD fellowship to N.K. (1SH6J24N).

## Declaration of competing interest

The authors declare the absence of competing interests.

## Author’s contributions

Conceptualization: S.P., H.D., S.A.

HyDrop bead design & generation: S.P., M.W.

Computational analysis: H.D., L.K., E.C.E., N.K.

Data collection, processing, and curation: H.D., F.D.R., G.H.

Experiments and sample preparation: K.T., F.D.R, V.C., H.D., N.R., K.S., R.V.

Resources: S.A., S.P.

Visualization: H.D., S.P., K.T.

Writing: H.D., S.P., K.T.

## Acknowledgments

We thank the members of the Laboratory of Computational Biology for their feedback during the process of method improvement and data collection. We thank Jean-Christophe Marine for his kind donation of the mouse melanoma lines. Lastly, we thank the staff of the Flemish Supercomputing Center (Vlaams Supercomputer Centrum - VSC) and VIB Data Core for their support.

## KEY RESOURCES TABLE

See supplementary material Table S1.

## STAR Methods

### EXPERIMENTAL MODEL AND STUDY PARTICIPANT DETAILS

#### Drosophila lines

The DGRP fly lines (DGRP-639, DGRP-502, DGRP-409) were purchased from Bloomington (25199, 28204, and 28278 respectively).

#### Mouse lines

Cortical brain tissue from female P60 with C57B/6J background was used. The experiments were conducted in line with KU Leuven’s ethical guidelines and approved by the Ethical Committee for Animal Experimentation (protocol number P007/2021).

#### Cell lines

For the barnyard test of HyDrop v2, cells from a human breast cancer cell line (MCF-7; RRID:CVCL_0031) and mouse melanoma cells were mixed. The use of the cell lines was approved by the KU Leuven Ethical Committee under project number S63316.

### METHOD DETAILS

#### Barcoded hydrogel bead manufacturing and storage

The microfluidic droplet generators for HyDrop v2 were produced following the previously described protocol^20^ modified as described below. A detailed protocol can be found at http://dx.doi.org/10.17504/protocols.io.8epv5n11dv1b/v1.

Dissolvable hydrogel beads for HyDrop v2 are synthesized similarly to previously published protocols^20,39^ and barcoded by three rounds of split-and-pool ligation reactions^40^. For synthesizing 2–3 mL batch of beads, 1.1 mL of Bead Monomer Mix (8.5-10% acrylamide (%T), 2-4% bisacryloylcystoylamine (%C), 10% Tris-buffered saline with EDTA and Triton X-100 (TBSET) (10 mM Tris-HCl pH 8, 137 mM NaCl, 2.7 mM KCl, 10 mM EDTA, 0.1% Triton X-100), 4 μM acrydite primer, 0.6% ammonium persulfate) was encapsulated into 60 μm diameter droplets in HFE-7500 Novac oil with EA-008 surfactant (RAN Biotech) with 0.5% TEMED. Flow rates: monomer: 400 and oil 750. The resulting emulsion was collected into 2 low bind 1.5 mL Eppendorf, layered with 200 μL mineral oil, and incubated at 65 °C for 14 hours, then paused at 12 °C. Excess mineral oil and the emulsion oil were removed and three washes with 1 mL of droplet-breaking solution (20% PFO in HFE) were performed. Beads were pelleted at 1000 xg, 4 °C for 60 seconds and washed three times in 1 mL of 1% SPAN-80 in hexane (0.2µm syringe filter). Beads were sequentially washed three times with TBSET and three times with TET buffer. Bead QC was performed as described previously (Zilionis et al., 2017; De Rop et al., 2022) and beads were filtered using a 70 μm filter (EASYstrainer from Greiner) to exclude large bead contamination. The beads are stored in TET buffer at 4 °C.

For barcoding round one, on the day of barcoding, the beads were first washed with TET buffer by spinning down (3 minutes at 1000 xg, gentle braking), discarding supernatant, and resuspension in TET buffer. After centrifugation, the TET buffer was removed, and the beads were washed two times with pre-ligation buffer (10 mM Tris-HCl pH. 8.0, 30 mM NaCl, 1 mM MgCl_2_ 0.1% Tween-20) for 1 minute at 1000 xg with gentle braking. After washing the supernatant was removed to obtain a compacted pellet of beads. A Hamilton microlab STAR robot was used for liquid handling in the deep well 96-well plates during barcoding. 22 µL of compacted beads was aliquoted in each well of the 96-well plate. Next, 16 µL of 2× ligation-primer buffer (100 mM Tris-HCl pH 7.5, 20 mM MgCl_2_) and 4 µL oligo mix cassette (200 µM) were added. The plate was spun, the contents mixed by vortexing, spun again, and placed on a PCR block (pre-heated to 75 °C with heated lid at 105 °C). To anneal the oligos the heating was turned off after 3 minutes at 75 °C which allowed gradual cooling to room temperature (about 2.5 hours). During this cooling process, the plate is taken off the PCR-block and vortexed every 30 minutes at 2000 RPM to resuspend the beads. When the temperature reached 33 °C the plate was left at room temperature for 10 minutes. The Mantis (Formulatrix) was used to aliquot 10 µL ligase mix (50 mM Tris-HCl pH 7.5, 10 mM MgCl2, 5.1 mM ATP, 100 U/µL T4 ligase (New England Biolabs)) to each well. Plates were sealed, vortexed in a vortex shaker and centrifuged for 30 seconds at 1000 xg. Then the beads were mixed on a thermoblock shaker for 30 seconds at 2000 RPM. Ligation was performed for 60 minutes at 25 °C with shaking alternating between 1000 and 1600 RPM for 30 seconds each. Afterwards, the plate was incubated overnight at 16 °C on a thermoblock while being shaked at 1000 RPM. After the overnight incubation, the plate was kept at 4 °C for at least 30 minutes before we proceeded with the inactivation of the ligation by performing a cleanup step on the Hamilton with STOP-25 (10 mM Tris-HCl pH 8, 25 mM EDTA, 0,10% Tween-20, 100 mM KCl). The beads were transferred to a 50 mL falcon and incubated for 20 minutes. Afterward, the beads were spun down (3220 xg for 7 minutes, gentle braking), the supernatant was removed, and the beads were transferred to a 15 mL tube. Next the beads were washed once with STOP-10 (10 mM Tris-HCl pH 8, 10 mM EDTA, 0.10% Tween-20, 100 mM KCl) and three times with TET buffer (centrifugation at 1000 xg for 1 minute). The second round of barcoding was started after two washes with pre-ligation buffer (centrifugation at 1000 xg for 1 minute) after which we transferred 22 µL compacted beads to each well of a 96-well plate.

For the round 2 barcoding, the 4 µL of the barcode 2 oligo cassette is added to 16 µL of 2× ligation-primer buffer in a 96-well plate and annealing is achieved again with a PCR-block. Afterwards, the plate is spun to collect the condensate. The annealed cassettes were then transferred to the bead plate. Ligation was performed as discussed in barcode step one.

Barcoding step 3 was done like step 1 with the exception that 23 µL of compacted beads was aliquoted in each well and mixed with 17 µL of 2× ligation-primer buffer and 3 µL of barcode 3 oligo mix (500 µM).

The barcoded beads are cleaned one last time on the Hamilton with STOP-25 buffer as described above and denatured with denaturation buffer (0.1 N NaOH, 1.68% Brij-35) for 10 min at room temperature at rotation (1 minute at 1000 xg). The beads are then washed with the neutralization buffer (100 mM Tris HCl pH 8, 10 mM EDTA, 0.1% Tween-20, 100 mM NaCl) for 10 min at room temperature with rotation (1 minute at 1000 xg). Finally, the beads are washed 3 times with TET buffer (1 minute at 1000 xg) and filtered twice using 70 um cell strainers.

The barcoded and cleaned beads are then incubated overnight at 4 °C in lysis buffer (125 mM Tris HCl pH 7, 150 mM NaCl, 12.5 mM MgCl_2_, 1.25% Triton X-100, 0.4% BSA). The beads are washed twice more with lysis buffer (1 minute at 1000 xg) and aliquoted into capped PCR-strip tubes (40 µL) and stored at –80 °C.

#### Sanger sequencing of barcoded beads

To test the purity of our barcoded HyDrop v2 beads, 1 µL of finalized barcoded beads was taken from their storage at −80 °C and they were pelleted and washed three times with 0.04% BSA in PBS in a 1.5 mL microcentrifuge tube (centrifuge for 1 minute at 500 xg). The beads were diluted with PBS/BSA under a stereo microscope (Leica S8 APO) on a petri dish (Falcon 351008) until individual beads could be picked with a Stripper® pipette and 75 µm capillaries (MXL3-STR and MXL3-75, CooperSurgical). Individual beads were transferred consequently over three drops (5 to 10 µL) to ensure no other beads were present in the capillary. Next, single beads, in 0.5 µL of PBS/BSA, were transferred to PCR tubes containing 1.5 µL PBS/BSA. Afterwards, the capillary was washed three times in a large volume of PBS/BSA.

To the picked beads in the PCR tube, 18 µL PCR master mix was added (final concentrations: 1× KAPA HiFi hot start ready mix (Roche Cat. No. KK2602), 30 mM DTT, 1 µM ATAC-QC oligo) and the following PCR program was used: 72 °C – 3 mins, 98 °C – 30 s, 22 cycles of 98 °C – 10 s; 59 °C – 30 s; 72 °C – 30 s; and finally the amplification products were cooled to 4 °C. This product was purified using 1.5× Ampure XP beads and the washed beads were resuspended in 10 µL of EB buffer (Qiagen) and 8 µL of the eluate was used in the next PCR reaction (total volume is 20 µL with final concentrations: 1× KAPA HiFi hot start ready mix, 1 µM P7 primer, 1 µM P5 primer). Sanger sequencing was done by LGC Genomics using the P7 primer.

To test HyDrop v1 bead purity the same protocol was followed but different oligos were used (see Table S1).

#### Husbandry and sample extraction

##### Mouse

The mice with C57B/6J background were housed under standard housing conditions in a pathogen-free facility with a 14 hr light, 10 hr dark light cycle from 7 to 21 hr. Mice used in the study were 60 days old. Animals were euthanized using 0.05 mL/g BodyWeight. The chest cavity was opened, and the brains were flushed with 5 mL PBS through the left ventricle of the heart followed by decapitation. Brains were dissected out and cortices were collected. The motor cortex is dissected, snap-frozen in liquid nitrogen, and stored at −80 °C.

##### Nuclei extraction

Mouse nuclei for HyDrop-ATACv2 and 10x Genomics Single Cell ATAC v2 were extracted from mouse cortex samples that were snap-frozen in liquid nitrogen and stored at −80 °C. We transferred ∼1 cm^3^ of frozen mouse cortex to 500 μL of ice-cold homogenization buffer (10 mM Tris-HCl pH 7.5, 10 mM NaCl, 3 mM MgCl_2_, 320 mM sucrose, 0.1 mM EDTA, 0.5% BSA, 0.1% IGEPAL CA-630, 1× complete protease inhibitor, and 1 mM DTT) in a Dounce Homogenizer (KIMBLE, 1 mL). The tissue was left to thaw for 2 min before the tissue was homogenized with 10 strokes of the loose pestle and 5 strokes of the tight pestle. Before the end of a 5-minute incubation, the homogenate was filtered through a 70 μm cell strainer (Corning), and the homogenizer and strainer were washed with homogenization buffer to obtain a final volume of 1 mL. The filtrate was transferred to a 1.5 mL DNA LoBind tube (Eppendorf) and centrifuged at 500 xg for 5 minutes. The supernatant was discarded, and the pellet was topped up to 520 μL with wash buffer 1 (10 mM Tris-HCl pH 7.5, 10 mM NaCl, 3 mM MgCl2, 320 mM sucrose, 0.1 mM EDTA, 0.5% BSA, 1× complete protease inhibitor, and 1 mM DTT) and resuspended. Next, we added 520 μL gradient medium (50% Optiprep, 10 mM Tris-HCl pH 7.5, 1 mM CaCl_2_, 5 mM MgCl_2_, 75 mM sucrose, 1× complete protease inhibitor, and 1 mM DTT) and mixed the entire volume by gentle pipetting. In a 2 mL DNA LoBind tube (Eppendorf) we layered 790 μL Optiprep cushion (29% Optiprep, 31 mM Tris-HCl pH 8, 77.5 mM KCl, 15.5 mM MgCl_2_, 129.2 mM sucrose) at the bottom and 1040 μL of the sample on top, gently without disrupting the cushion. The tube was then centrifuged at 9000 xg for 20 minutes at 4 °C. After the centrifugation the debris on top was carefully removed with a 1 mL tip and the rest of the supernatant was also carefully removed until 50 μL was left in the tube. The pellet was gently resuspended before we mixed it with 50 μL 2× permeabilization buffer (20 mM Tris-HCl pH 7.5, 20 mM NaCl, 6 mM MgCl_2_, 0.2% Tween-20, 0.02% IGEPAL CA-630, 0.02% Digitonin, 2% BSA, and 2 mM DTT) and left the nuclei suspension for 2 minutes on ice before we stopped the permeabilization by mixing it with 1 mL of wash buffer 2 (10 mM Tris-HCl pH 7.5, 10 mM NaCl, 3 mM MgCl_2_, 0.1% Tween-20, 1% BSA). The nuclei were pelleted by centrifugation at 500 xg for 5 minutes at 4 °C. The nuclei were resuspended in PBS with 0.04% BSA for HyDrop-ATAC v2 and in Diluted Nuclei Buffer for 10x Genomics Single Cell ATAC v2 processing.

##### Fly

All flies were raised on a yeast-based medium and kept at 25 °C on a 12h/12h day/night light cycle. For embryo collection, flies were transferred to cages fixed to a plate with a normal yeast-based medium 24 hours before embryo collection. On the day of embryo collection, the plate was exchanged for a juice plate. The juice was prepped by adding 20 g agar and 22 g sucrose to 300 ml distilled water, boiling, and then mixing in 100 ml apple juice, 10 ml 95% ethanol, and 5 ml 100% glacial acetic acid. The flies laid eggs for four hours. Sixteen hours later, the embryos aged between 16 to 20 hours were collected from the juice plate with a small brush and transferred to a 70 µm nylon mesh filter. To dechorionate the embryos, the embryos were treated for 2 min with 5% bleach and then washed 4 times with 0.1% Triton X-100 in 1× PBS. After a final wash in 1× PBS, the embryos were transferred to a 1.5 ml tube, spun down for 2 min at 500 g, supernatant was removed, and the embryo pellet was snap frozen on dry ice and stored at −80°C.

##### Nuclei extraction

Nuclei were isolated by using an adapted protocol based on the nuclei isolation protocol for single-cell ATAC sequencing (10x Chromium CG000169-RevE). Briefly, the embryos were resuspended in 500 µl cold lysis buffer (10 mM Tris-HCl pH 7.4, 10 mM NaCl, 3 mM MgCl2, 0.1% Tween-20, 0.1% IGEPAL CA-630, 0.01% Digitonin, 1% BSA), transferred to a Dounce homogenizer and incubated on ice for 5 minutes. The tissue was disrupted by 25 strokes with the loose pestle, incubated on ice for 10 minutes, and disrupted by 25 strokes with the tight pestle. To remove debris, the solution was filtered on a 10 µm nylon mesh filter and washed with 1 ml wash buffer (10 mM Tris-HCl pH 7.4, 10 mM NaCl, 3 mM MgCl_2_, 0.1% Tween-20, 1% BSA). Nuclei were collected by centrifugation at 500 g for 5 mins at 4°C, supernatants were carefully removed, and nuclei were resuspended in 50 µl 1× Diluted Nuclei Buffer (10x Chromium). Nuclei quality and concentration were assessed by the LUNA-FL Dual Fluorescence Cell Counter.

##### Cell lines

For a detailed description of cell culture and cell dissociation see De Rop (2022). In brief, MCF-7 cells (NCI-DTP Cat# MCF7, RRID:CVCL_0031) were cultured in RPMI1640 (ThermoFisher 11875093) medium supplemented with 10% FBS (ThermoFisher 10270–106), 1% penicillin/streptomycin (Life Technologies 15140122), and 10 ug/mL insulin (Sigma Aldrich I9278) and passaged twice per week. Mouse melanoma cells were cultured in DMEM (ThermoFisher 13345364) supplemented with 10% FBS and 1% penicillin/streptomycin and passaged once per week. All cell lines tested negative for mycoplasma prior to use. Cells were washed in PBS and dissociated into single-cell suspensions by adding 1.5 mL of 0.05% trypsin (Life Technologies 25300054) and waiting for 5 minutes. The single-cell suspension was centrifuged at 500 rcf for 5 min at 4°C and the resulting pellet was resuspended in PBS. This PBS wash was repeated once more and the single-cell suspension was processed further.

##### Nuclei extraction

For the barnyard, the cell lines were mixed to equal parts and a pellet of one million dissociated cells or fewer was incubated on ice in 200 μL of ATAC lysis buffer (1% BSA, 10 mM Tris-HCl pH 7.5, 10 mM NaCl, 0.1% Tween-20, 0.1% NP-40, 3 mM MgCl_2_, 70 μM Pitstop in DMSO, 0.01% digitonin) for 5 minutes. 1 mL of ATAC nuclei wash buffer (1% BSA, 10 mM Tris-HCl pH 7.5, 0.1% Tween-20, 10 mM NaCl, 3 mM MgCl_2_) was added and the nuclei were pelleted at 500 xg, 4 °C for 5 minutes. The resulting pellet was resuspended in 100 μL of ice-cold PBS and filtered with a 40 μm strainer (Flowmi).

#### HyDrop-ATAC v2 library preparation

A detailed description of the protocol can be found here dx.doi.org/10.17504/protocols.io.x54v97mmpg3e/v1.

##### Mouse

Between 7500 and 18000 nuclei, resuspended in PBS (supplemented with 0.04% BSA), were tagmented for one hour at 37 °C in 15 μL ATAC reaction mix (10% DMF, 10 mM Tris-HCl pH 7.4, 5 mM MgCl_2_, 5 ng/µL Tn5, 70 µM Pitstop in DMSO, 0.1% Tween-20, 0.01% Digitonin). Tagmented nuclei were placed on ice and 95 µL of PCR mix (1.25× Phusion HF buffer, 15% Optiprep, 1.25 mM dNTPs, 75 mM DTT, 0.0625 U/µL Phusion HF polymerase, 0.0625 U/μL Deep Vent polymerase, 0.0125 U/μL ET SSB) was added. Next, 110 μL of nuclei mixed in PCR mix was co-encapsulated with 45 μL of HyDrop-ATACv2 beads in HFE-7500 Novac oil with EA-008 surfactant (RAN Biotech) using the Onyx microfluidics platform (Droplet Genomics). The resulting emulsion was split over two PCR tubes and was placed in a thermocycler with the following program: 72 °C – 15 mins, 98 °C – 3 mins, 12 cycles of 98 °C – 10 s; 55 °C – 30 s; 72 °C – 1 min; and a final extension of 72 °C – 5 min, followed by a hold stage at 4 °C.. To break the emulsion and remove excess oil, 125 μL of recovery agent (20% PFO in HFE) was added and the PCR tubes were inverted 10 times to mix before the oily phase was removed. Next, we added 180 μL GITC mix (5M GITC, 25 mM EDTA, 50 mM Tris-HCl pH 7.4), 10 μL Dynabeads, and 10 μL 2.4 M DTT to each of the aliquots, and mixed everything by pipetting up and down 10 times and incubating for 10 minutes. Dynabeads were pelleted on a magnet and washed twice with 80% ethanol. Elution was done in 50 μL of EB-DTT-Tween (24 mM DTT, 0.1% Tween-20 in EB (10 mM Tris-HCl pH 8.5)). Next, a 1.2× Ampure bead purification was performed according to manufacturer’s recommendations. Elution was done with 40 μL of EB-DTT (10 mM DTT in EB). The library was completed by amplifying the eluate in a total volume of 100 μL PCR mix (1× KAPA HiFi, 1 μM index i7 primer, 1 μM Universal P5 primer) with the following program: 95 °C – 3 mins, 12 cycles of 98 °C – 10 s; 63°C – 30 s; 72 °C – 1 min; and a final extension of 72 °C – 1 min, followed by a hold stage at 4 °C. The final libraries were subjected to 0.4×–1.2× doublesided Ampure purification and eluted in 20 μL elution buffer (Qiagen).

##### Fly

Analogous to mouse nuclei, the HyDrop v2 protocol was performed on the lysed Drosophila nuclei, resuspended in 1x nuclei buffer. A total of 80,000 nuclei were resuspended in 40 μL of ATAC reaction mix (10% DMF, 10% Tris-HCl pH 7.5, 5 mM MgCl2, 4.7 ng/μL Tn5, 70 μM Pitstop in DMSO, 0.1% Tween-20, 0.01% digitonin) and incubated at 37°C for 1 hr. Tagmented nuclei were placed on ice and counted with the LUNA-FL Dual Fluorescence Cell Counter. Per HyDrop reaction, a total of 37,500 tagmented nuclei were used in a 80 µl PCR reaction (1.37× Phusion HF buffer, 1.37% PEG-8000, 0.68 mM dNTPs, 89 mM DTT, 0.04 U/µL Phusion HF polymerase, 0.04 U/μL Deep Vent polymerase, 0.01 U/μL ET SSB). Next, 80 μL of nuclei mixed in PCR mix was co-encapsulated with 35 μL of HyDrop-ATACv2 beads in HFE-7500 Novac oil with EA-008 surfactant (RAN Biotech) using the Onyx microfluidics platform (Droplet Genomics). The resulting emulsion was placed in a thermocycler with the following program: 72 °C – 20 mins, 98 °C – 3 mins, 12 cycles of 98 °C – 10 s; 58 °C – 30 s; 72 °C – 1 min; and a final extension of 72 °C – 5 min, followed by a hold stage at 4 °C. To break the emulsion and remove excess oil, 125 μL of recovery agent (20% PFO in HFE) was added and the PCR tube was inverted 10 times to mix before the oily phase was removed. Next, we added 180 μL GITC mix (5M GITC, 25 mM EDTA, 50 mM Tris-HCl pH 7.4), 10 μL Dynabeads Silane One, and 10 μL 2.4 M DTT, and mixed everything by pipetting up and down 10 times and incubating for 10 minutes. Dynabeads were pelleted on a magnet and washed twice with 80% ethanol. Elution was done in 100 μL of EB-DTT-Tween (24 mM DTT, 0.1% Tween-20 in EB (10 mM Tris-HCl pH 8.5)). Next, a 1.1× Ampure bead purification was performed according to manufacturer’s recommendations. Elution was done with 40 μL of EB-DTT (10 mM DTT in EB). The library was completed by amplifying the eluate in a total volume of 100 μL PCR mix (1× KAPA HiFi, 1 μM index i7 primer, 1 μM Universal P5 primer) with the following program: 95 °C – 3 mins, 10 cycles of 98 °C – 10 s; 63°C – 30 s; 72 °C – 1 min; and a final extension of 72 °C – 1 min, followed by a hold stage at 4 °C. The final libraries were subjected to 0.4×–1.2× doublesided Ampure purification and eluted in 20 μL elution buffer (Qiagen).

scATAC-seq libraries were prepared according to the Chromium Single Cell ATAC reagent kits v2 user guide (10x Genomics, CG000496 Rev B). Briefly, the transposition reaction was prepared by mixing the desired number of nuclei with ATAC Buffer and ATAC Enzyme, and was then incubated for 30 minutes at 37 °C. Tagmented nuclei were partitioned into nanoliter-scale gel bead-in-emulsions (GEMs). DNA linear amplification was then performed by incubating the GEMs under the following thermal cycling conditions: 72 °C – 5 min, 98 °C – 30 s, 12 cycles of 98 °C – 10 s; 59 °C – 30 s; 72 °C – 1 min, and finally a hold stage at 4 °C. GEMs were broken using Recovery Agent, and the resulting DNA was purified by sequential Dynabeads and SPRIselect reagent beads cleanups. Libraries were indexed by PCR using a Single Index kit N set A and incubated under the following thermal cycling conditions: 98 °C – 45 s, seven cycles of 98 °C – 20 s; 67 °C – 30 s; 72 °C – 20 s; and a final extension of 72 °C – 1 min, followed by a hold stage at 4 °C. Sequencing libraries were subjected to a final bead cleanup with SPRIselect reagent.

#### Sequencing

HyDrop-ATAC v2 libraries were sequenced on an Illumina NextSeq2000 or NovaSeqX system using at least 43 cycles for read 1 (ATAC paired-end mate 1), 10 cycles for index 1 (sample index), 41 cycles for index 2 (HyDrop-ATAC v2 barcode), and 44 cycles for read 2 (ATAC paired-end mate 2).

#### scATAC data processing

All in-house generated 10x and HyDrop samples were processed using the PUMATAC pipeline^19^ (RRID:SCR_026624), built on Nextflow^41^ (v21.04.3; RRID:SCR_024135). With PUMATAC, we aligned the samples to the mm10 and dm6 reference genomes for mouse and fly respectively to write fragment files and perform quality assessment. For an in-depth description of the pipeline, see De Rop et al. (2023)^19^ and https://github.com/aertslab/PUMATAC.

##### Barnyard

A mixed library of 5000 mouse melanoma and 5000 MCF-7 cells was generated following the standard HyDrop v2 library protocol as described above for the mouse cortex. The raw data was processed using the PUMATAC pipeline^19^ and aligned to a combined reference genome for mm10 and GRCh38. Cut-offs for TSS enrichment and number of unique fragments per cell were set using the Otsu algorithm. Lastly, the fragments belonging to each genome were counted and compared.

##### Mouse samples

10x v2 and 10x v1 fastq files were downloaded from 10x Genomics database. The samples were processed using the PUMATAC pipeline^19^, together with the in-house generated data. Cut-offs were set per sample using the Otsu threshold for TSS enrichment and number of unique fragments per cell. Next, fragments overlapping earlier published mouse candidate cis-regulatory regions^24^ were combined into a count matrix using pycisTopic^5^ (v2.0a0, RRID: SCR_026618). Potential doublets were removed using Scrublet (RRID:SCR_018098) with an automatic threshold set based on the expectation of 10% doublets. Cells were clustered based on initial pycisTopic^5^ Latent Dirichlet Allocation (LDA) using 80 topics. Based on this initial clustering, a new set of consensus regions was generated using MACS2, overlapping fragments recounted, and assembled into a second count matrix. Again, doublets were removed as previously and LDA performed with now 130 topics to capture all possible cell types. Artifacts attributed to batch effects were corrected using Harmony^42^ giving the sequencing technique as variable instead of the sample identifier. Next, based on Leiden resolution 2.5, the cells were annotated using differentially accessible regions (DARs) of previously published datasets^25,43^. The newly annotated cell types were subdivided based on the sequencing technique of each sample. On the new variable, DARs were calculated based on imputed chromatin accessibility. Here, only regions passing one-versus-all Wilcoxon rank-sum tests with 0.05 adjusted p-value are included. The top 1000 DARs per cell type per sequencing technique were investigated with motif enrichment analysis with pyCisTarget^10^ (v1.0a2, RRID:SCR_026626). Using pycisTopic^5^ to call peaks per cell type in pseudobulk with MACS2, peak BED files per cell type were generated.

For BigWig files generation per cell type/sequencing technique, the lowest number of cells in one cell type in the respective sparsest sequencing technique was set as maximum. An equal number of cells for that cell type was randomly selected per remaining sequencing technique. Only cell types with more than 100 cells in each sequencing technique were included further on. Next, BigWig files for fragment coverage and Tn5 cut sites were generated for the downsampled, equal number of cells per technique with scATAC Fragment Tools (<u>RRID:SCR_026643</u>) normalizing the genome coverage (divided by the number of fragments. They were visualized with the integrative genome viewer (IGV, RRID:SCR_011793)^44^. The carrot plots (i.e., heat maps) were generated with deepTools2^45^ (v3.5.3; RRID:SCR_016366) plotting the accessibility per cell type and sequencing technique (BigWig) on the cell type-specific peak files (i.e., all regions accessible in that respective cell type). This was done for BigWigs based on fragment coverage and Tn5 cut sites.

For the comparison of quality metrics across microfluidic techniques, the samples were downsampled to 36,000 reads per cell (RPC) as indicated in the respective figures. The plots were generated based on PUMATAC quality control evaluation with cell names defined based on previous annotations of the full sequencing depth count matrix.

##### Drosophila samples

Analogous to the mouse cortex data processing, the 10x v2 and HyDrop v2 data were processed with the PUMATAC pipeline^19^. Cut-offs were set per sample using the Otsu threshold with a maximum cut-off of 800 unique fragments and a TSS enrichment of 2. Fragments overlapping candidate regions of the earlier published Drosophila fly embryo dataset^27^ were combined into a count matrix using pycisTopic^5^ and clustered with 120 topics. Based on this initial clustering, a new set of consensus regions was generated with MACS2, overlapping fragments recounted and assembled into a new count matrix. LDA was performed in pycisTopic^5^ with 220 topics and clusters annotated with DARs from the published Drosophila embryo atlas^27^ on Leiden resolution 1.2. Correction of batch effects was performed with Harmony^42^ using the respective wet lab technique (sequencing technique and sample preparation parameter) as variable. Analogous to the mouse data, the annotated cell types were subdivided among HyDrop v2 and 10x v2. On the new variable, DARs were calculated based on imputed chromatin accessibility. Here, only regions passing one-versus-all Wilcoxon rank-sum tests with 0.05 adjusted p-value are included. The top 1000 DARs per cell type per sequencing technique were investigated with motif enrichment analysis with pyCisTarget^8^. Using pycisTopic^3^ to call peaks per cell type in pseudobulk with MACS2, peak BED files per cell type were generated.

For BigWig files generation per cell type/sequencing technique, the lowest number of cells in one cell type in the respective sparsest sequencing technique was set as maximum. An equal number of cells for that cell type was randomly selected per remaining sequencing technique. Only cell types with more than 1000 cells in each sequencing technique were included further on. Next, BigWig files for fragment coverage and Tn5 cut sites were generated for the downsampled, equal number of cells per technique with scATAC Fragment Tools (https://aertslab.github.io/scatac_fragment_tools/) normalizing the genome coverage (divided by the number of fragments. They were visualized with the integrative genome viewer (IGV; RRID:SCR_011793)^44^ Just as in mouse, the carrot plots (i.e., heat maps) were generated with deepTools2^45^ for BigWigs based on fragment coverage and Tn5 cut sites.

For the comparison of quality metrics between HyDrop v2 and 10x v2, the samples were downsampled to 12,000 reads per cell (RPC) as indicated in the respective figures. The plots were generated based on PUMATAC quality control evaluation with cell names defined based on previous annotations of the full sequencing depth count matrix.

For the sciATAC data, the available fragment files of two experiments of embryos of 16 to 20 hours after egg laying form the publicly available data from the embryo atlas^27^ were downloaded. Just as previously described, they were scored in two rounds with their own consensus peaks in the second processing round. Here, the maximum cutoff was just as in 10x and HyDrop samples set to TSS enrichment of two and 800 unique fragments. The data is clustered using pycisTopic^5^ with 220 topics and previous annotations^27^ assigned. Again, DARs were calculated as previously described, and motif enrichment analysis was performed with pyCisTarget^8^ on the top 1000 DARs. Coverage and cut site BigWig files were calculated analogous to 10xv2 and HyDrop v2 fly data. No downsampling took place for sciATAC data.

#### Model training

##### Mouse

Samples generated with the HyDrop v2 platform were extracted from the count matrix generated for the integrated samples. Again, using pycisTopic^5^, LDA was performed with 100 topics. No batch correction of the HyDrop v2 beads was needed. For downstream training of the HyDrop v2-based sequence model, pseudobulk profiles were generated in pycisTopic^5^ with MACS2 peak calling based on grouping cells in motor cortex cell types previously annotations of the full data set.

The sequence model was trained with the CREsted package^22^ (v.1.3.0RRID:SCR_026617) following the default peak regression parameters. Briefly, the data was divided into validation (chr8, chr10), train (chr 9, chr18), and remaining data and resized to 2,114bp length. Peaks were normalized using the top 2% of the peaks. Next, the data was augmented with a sequence shift of 3 bp (to both sites) stochastically during training and the use of the reverse complement previous to model training using the dilatedcnn model architecture.

To compare the HyDrop v2 sequence model to a published and evaluated data set, 10x multiome data generated with frozen primary motor cortex tissue from eight P56 C57BL/6J male mice (*Mus musculus*) generated^46^ with the 10x protocol was used. The dataset has previously been annotated and extensively evaluated for deep learning purposes^12,46^. The model was trained using the standard CREsted peak regression pipeline, by first pretraining on all consensus peaks and fine-tuning on a set of cell type-specific peaks. Just as in the HyDrop v2 model training, the same validation and training splits were handled followed by resizing to 2,114bp length. The peaks were also normalized using the top 3% of the peaks and the data augments as previously described. Again, the dilatedcnn model architecture was used for the model training.

In both models, recurrent sequence patterns were identified in the top 2,000 regions that overlapping between 10x and HyDrop v2 data sets per cell type with *tfmodisco-lite* within the default parameters of the CREsted package.

Patterns between cell types and techniques were compared with the function *crested.tl.modisco.process_patterns* based on tomtom with the default parameters for similarity threshold of 3.5, trimming patterns based on an information content of 0.1 and discarding patterns with a threshold of 0.2.

###### *In vivo* validated enhancers recovery curves

A set of 122 functionally validated On-Target, mouse genomic enhancer regions in the mouse motor cortex was taken from a dataset provided by Ben-Simon et al.^25^. We evaluated the HyDrop v2 and 10x mouse motor cortex models by taking their predictions for these regions and calculating specificity through the *crested.pp._utils._calc_proportion* function. The specificity scores per model were then used to assess precision and recall, as was done in previous work^12^. Similarly, from the scATAC-seq peak heights, we calculated specificity scores to obtain precision and recall scores. Ground truth labels were obtained from the annotation of the validated enhancers.

As the 10x dataset had three cell types not found in the HyDrop dataset (SstChodl, Sncg, and VLMC), we merged them into the Sst, Vip and Endo classes respectively by taking the max prediction and peak heights between the merged classes. The HyDrop v2 dataset on the other hand has a L4IT class not present in the 10x data, which is why it was merged with the L5IT class also based on maximal prediction and peak height between the two classes.

##### Fly

Samples generated with the HyDrop v2 platform and 10x v2 were extracted from the count matrix generated for the integrated samples into two count matrixes. Again, using pycisTopic^5^, LDA was performed with 220 topics for HyDrop v2 and 200 topics for 10x v2. For downstream training of sequence models per sequencing technique, pseudobulk profiles were generated in pycisTopic^5^ with MACS2 peak calling based on grouping cells in previously annotated cell types from the full data set excluding unannotated clusters.

The S2F model was trained with the CREsted package following a version of the default peak regression parameters adapted for fly. Briefly, the data was divided into train, validation and test sets. The regions in chromosome 2R were evenly divided into two to use as validation and test sets and the remaining chromosomes were used as the training set. The original region length of 500bp was kept. Peak heights were normalized per chromosome to a target mean accessibility of 0.5. After normalization, z-scores of peak heights were calculated per region. For each cell type, top 3000 regions with the highest z-scores were kept and the accessibility values of all the other regions were set to zero for that cell type. CosineMSELoss (from crested.tl.losses) was used along with the default optimizer and metrics from default CREsted peak regression configuration. The *deeptopic_cnn* model architecture was used with the following optional parameters: *filters=500, conv_do=0.5.* Since the default *deeptopic_cnn* model architecture has an output activation that’s incompatible with regression task, it was replaced with a *softplus* activation. Two forms of augmentations were used: Before training, the training data was augmented using reverse complementation, this is achieved by adding the reverse complement of each sequence to the training data keeping the same target vector as the original sequence. During the training, each input region was stochastically shifted up to 3bp on either direction, still keeping the original target vector.

We identified the top 2,000 highest predicted regions per cell type per model and calculated the contribution scores of regions that are in the intersection of HyDrop v2 and 10x v2 data sets for the respective cell type. Later, these input sequences and contribution scores were used as input to *tfmodisco lite*-software (RRID:SCR_024811), using the available wrappers in CREsted package to find the most important patterns identified by 10x and HyDrop v2 models per cell type. An importance threshold of 5 was handled to select the most important motifs displayed

###### 10-fold cross-validation and selection of 500 bp regions in 2kb window

To evaluate the ability of our S2F models to capture biologically relevant information, we made use of validated enhancers downloaded from https://enhancers.starklab.org/ and lifted over to dm6 genome annotation with UCSC. The original cell type annotations were manually translated into the cell types used in this paper (Table S2). Each ∼2kb enhancer region (merged for the annotated cell types) was scanned with a sliding window approach of 500 bp windows in steps of 10bp over the whole enhancer with the 10-fold cross-validated models. Then, the 500bp regions showing the highest CREsted predicted accessibility for all matching annotated cell types were selected based on the highest correlations between predicted and ground truth insertions.

###### sciATAC-seq based models

The trained models were obtained from https://doi.org/10.5281/zenodo.8011697. The enhancer activity models for used for all the analysis. The 501bp synthetic DNA sequences were acquired from Almedia et al.^15^. The sequences were flanked on each side by 250bp random sequences to acquire a 1001bp sequence. The nucleotide contribution scores were calculated for one replicate model for each of the ten cross-validation folds, from the relevant tissue (except for the epidermis, where none of the provided replicates of fold 5 were working). The final contribution score was calculated by averaging all the contribution scores per sequence. The visualization of the contribution scores was done in CREsted software.

#### Footprinting with PRINT

Footprint scores calculations were carried out using the default PRINT v3 procedure as described in https://ruochiz.com/scprinter_doc/index.html (scPRINTER python implementation of PRINT, https://zenodo.org/records/13963610). For both, mouse and fly, we used the estimated Tn5 bias files provided by the original authors. As Tn5 insertion inputs, we used the Hydrop v2 and 10x v2 fragment files preprocessed by PUMATAC pipeline^19^ for in-house generated data and cellranger-arc (10x Genomics) for public the mouse multiome cortex data^46^. The data was grouped based on the cell barcodes annotated as previously described.

For mouse, we inspected the footprints of validated enhancer regions from the BICCN challenge^12^. For fly, we calculated footprints for all Drosophila embryo enhancers^17^ active in stages 15-16 downloaded from https://enhancers.starklab.org/ and lifted over to dm6 genome annotation with UCSC. Each ∼2kb enhancer region was scanned with a sliding window approach of 500 bp windows in steps of 10bp over the whole enhancer. Then, the 500bp regions showing the highest CREsted predicted accessibility for all matching annotated cell types were selected for further analysis based on the highest correlations between predicted and ground truth insertions.

## Supplementary material

**Fig S1.**
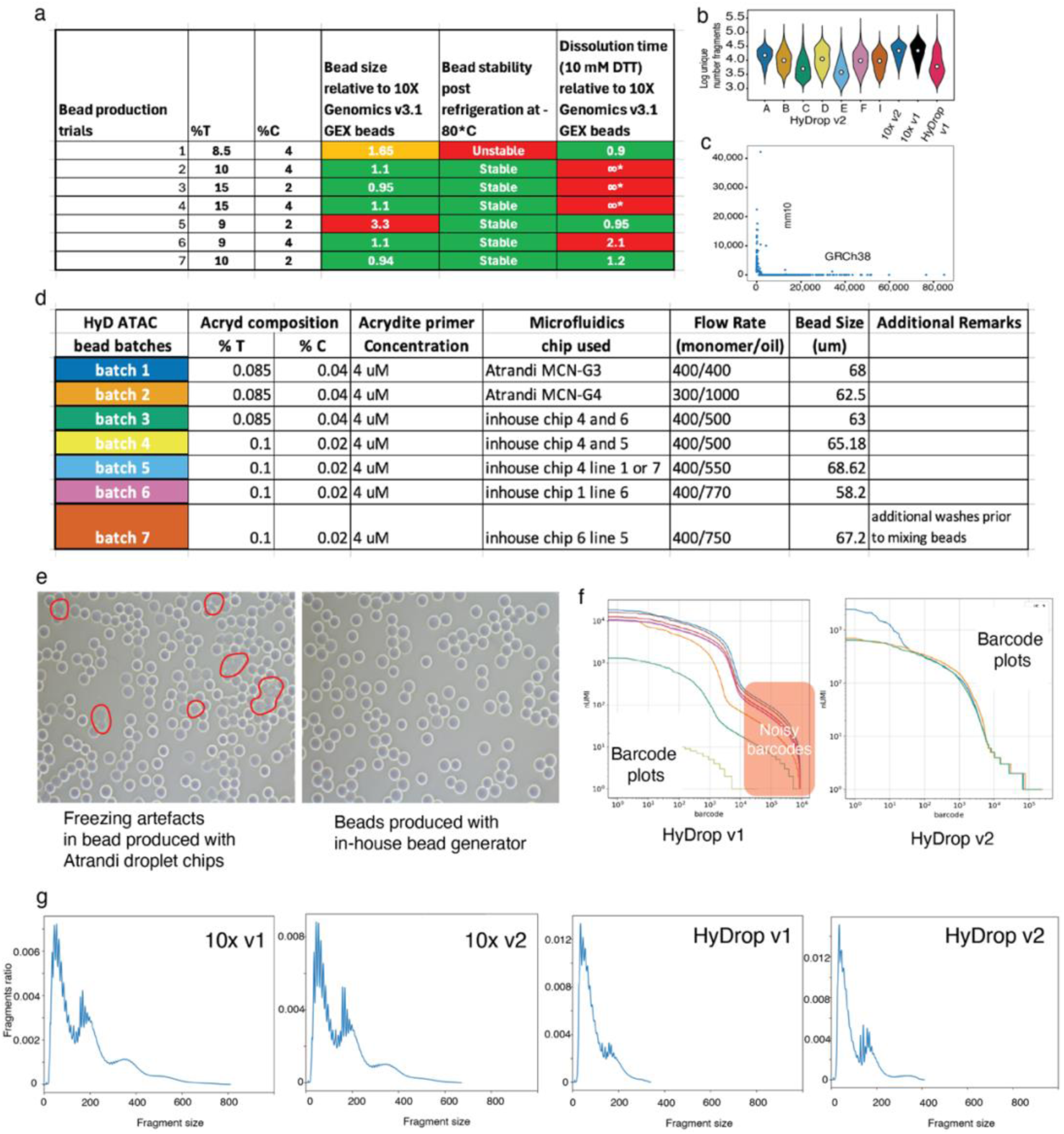
Improvement of HyDrop v2 bead generation. **a**, Characterization of bead quality and stability with different monomer compositions. Different percentages of total monomer and cross linker composition is varied during different bead productions and the stability of the beads are evaluated using three different characteristics: i.) bead size variations, ii.) bead stability post refrigeration, and iii.) dissolution time of the beads in a reducing environment relative to 10X genomics NextGEM 3’ RNA beads. **b,** log-transformed number of unique fragments of experiments pooled for seven different bead batches of HyDrop v2 compared to pooled 10x v2, 10xv1, and HyDrop v1 samples. The median is shown as white dot. **c**, Barnyard of HyDrop v2 protocol with mouse and human cell line. **d.** Table illustrating different parameter and characteristics of different bead production batches. **e.** Bright field image of the beads showing freezing artifact in suboptimal bead batch productions. **f.** Barcode ranking plot showing noisy barcode contamination in the HyDrop v1 beads arising from the barcode bleeding during the production. **g**, Fragment size plot of 10x v1, 10x v2, HyDrop v1, and HyDrop v2 experiment. 10x data shown is downloaded from 10x Genomics website.

**Fig. S2.**
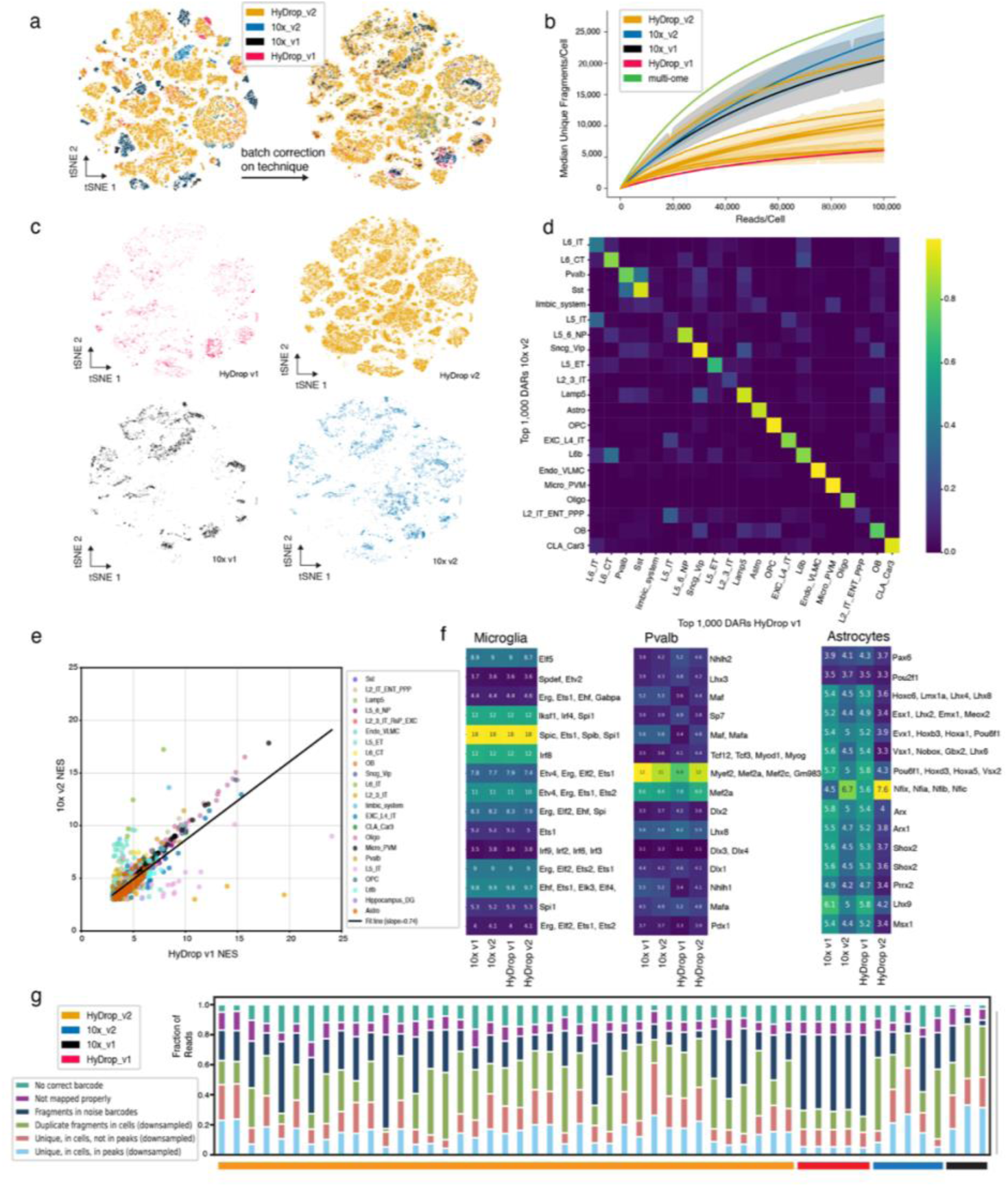
Quality metrics across techniques in mouse cortex data. **a**, *t*-distributed stochastic neighbor embedding (tSNE) of all 141,010 cells across 44 experiments colored by technique used, left: no batch correction, right: batch corrected for the used technique. Data points were randomly shuffled before plotting. **b**, Sequencing efficiency compared across 10x v1 and v2, multi-ome (black), HyDrop v1 and HyDrop v2 beads (7 batches). The colors are shown in a. **c**, *t*-distributed stochastic neighbor embedding (tSNE) showing the distribution of cells in combined embedding in a, 2 experiments 10x v1 with a total of 10,177 cells, 2 experiments 10x v2 with a total of 13,123 cells, 5 experiments HyDrop v1 with a total of 7,163 cells, 35 experiments HyDrop v2 with a total of 110,547 cells). **d**, Heatmap for top 1000 DARs per cell type correlated between 10x v2 and HyDrop v1 data. **e**, Scatterplot of normalized enrichment score (NES) of cell-type-specific transcription motifs, dots represent a common motif between HyDrop v1 and 10x v2 data colored per cell type. **f**, NES of microglia, Pvalb, and astrocytes divided per technique. **g**, Stacked bar plot showing the fraction of reads across each step of data processing using the PUMATAC pipeline (‘Unique, in cells, in peaks’: final fraction of sequencing reads retained in count matrices). The colors at the bottom of the plot indicate the technique of the experiment. The data is down-sampled to 36kRPC.

**Figure S3.**
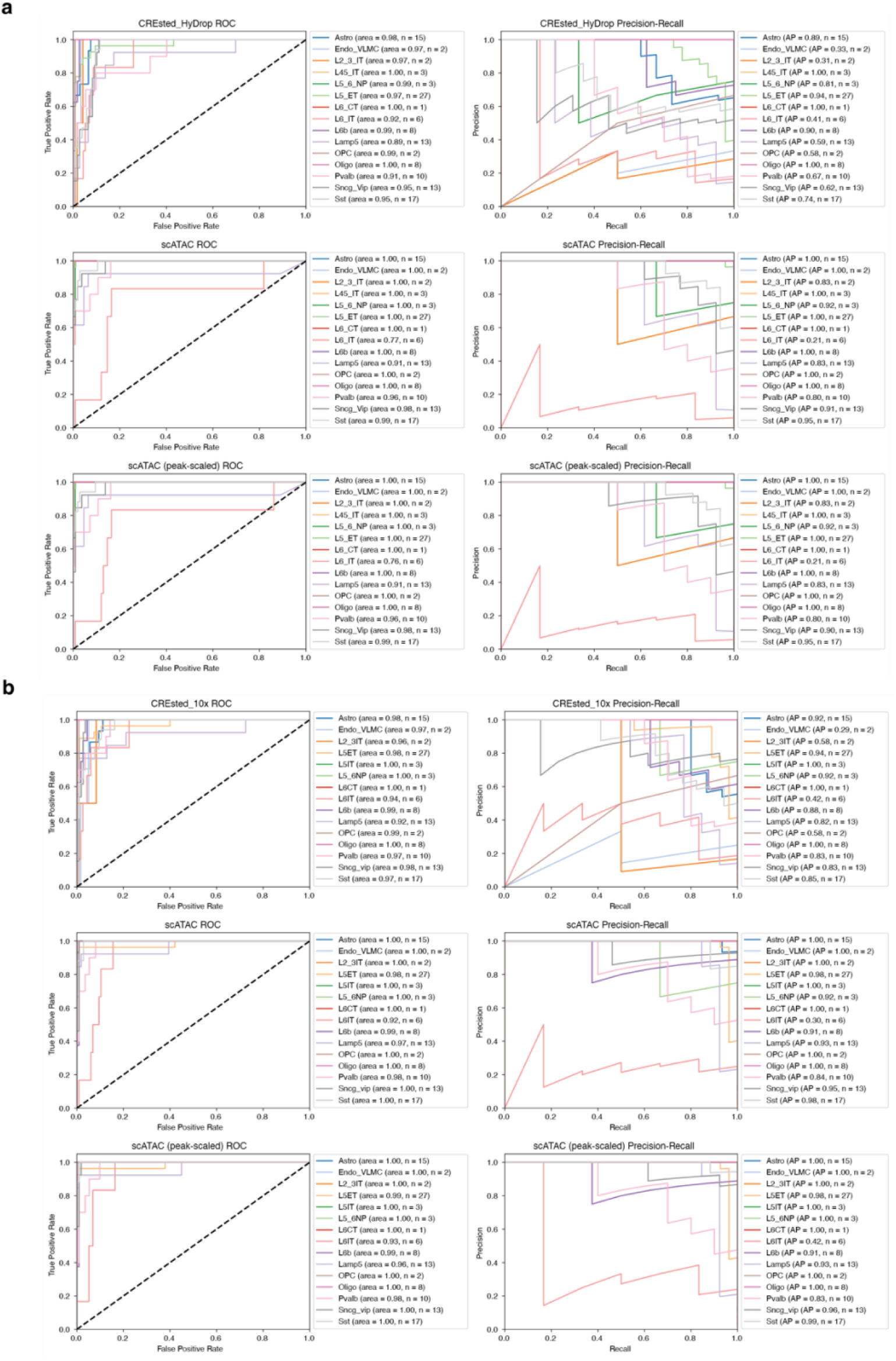
Overview of prediction performance on *in vivo* validated mouse motor cortex enhancers per cell type. ROC and PR curves from specificity scores per cell type on *in vivo* validated of the Hydrop v2 CREsted model, HyDrop scATAC-seq data and peak-scaled HyDrop v2 scATAC-seq data (**a)**, and of the 10x CREsted model, 10x scATAC-seq data and peak-scaled 10x scATAC-seq data (**b**).

**Fig. S4.**
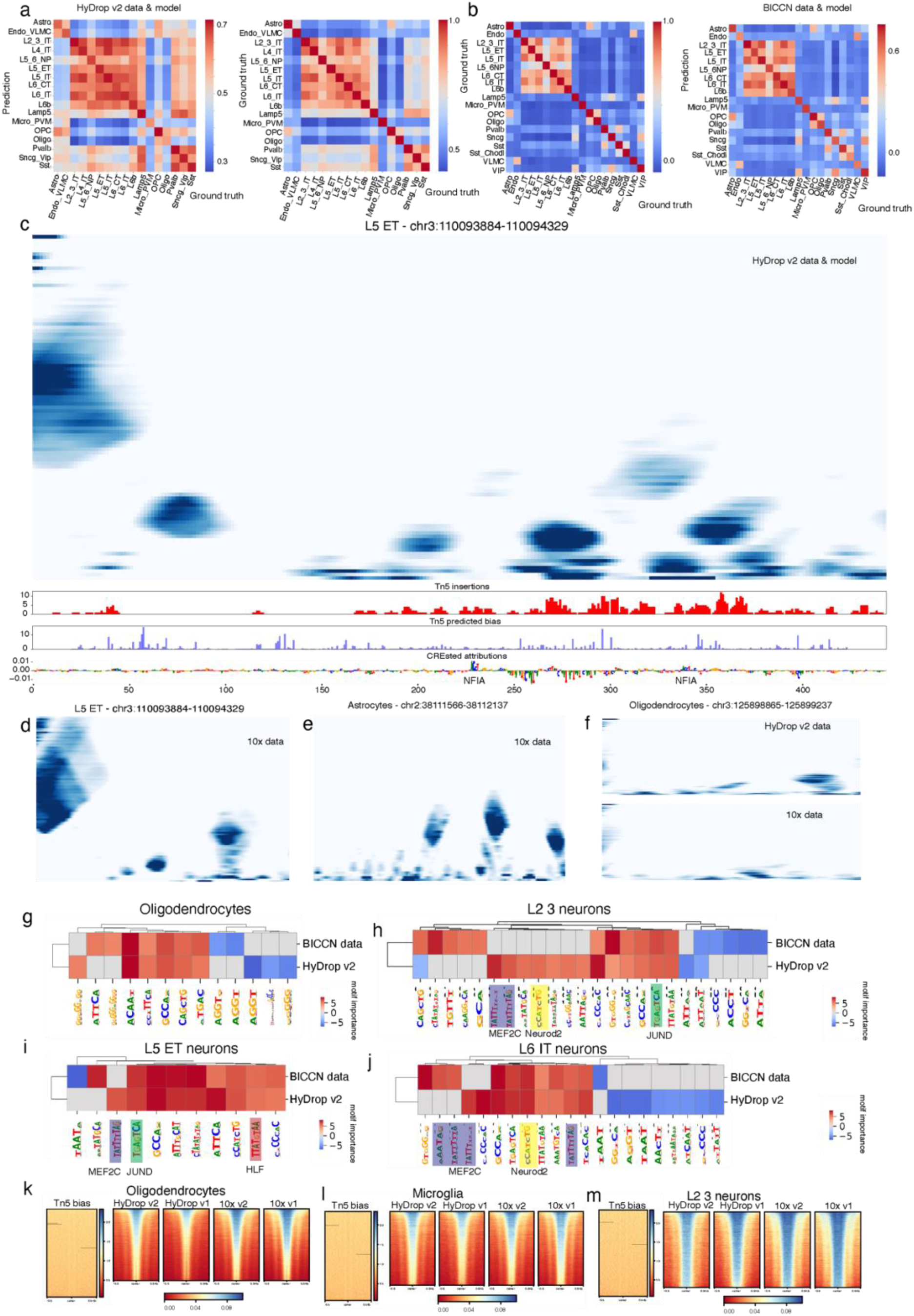
Sequence model comparison and multiscale footprinting in mouse cortex. **a**, Heatmap of correlation of prediction by HyDrop v2 model with ground truth (left), self-correlation of HyDrop v2 data as ground truth (right) for HyDrop v2 data. **b**, Heatmap of correlation of prediction by 10x v2 model with ground truth (right), self-correlation of 10x v2 data as ground truth (left) for 10x v2 data. **c**, Multiscale footprint of the region (mm10 chr3:110093884-110094329) in L5 ET neurons from HyDrop v2 data. Bottom tracks show Tn5 insertion, the predicted Tn5 bias and the nucleotide contribution scores based on the HyDrop v2 sequence model. Previously described TF binding sites are highlighted for the corresponding TF. **d**, Multiscale footprint of the region (mm10 chr3:110093884-110094329) in L5 ET neurons from 10x data. **e,** Multiscale footprint of the region (mm10 chr3:110093884-110094329) in astrocytes from 10x data (HyDrop v2 shown in Figure 3j. **f**, Multiscale footprint of the region (mm10 chr3:125898865-125899237) in oligodendrocytes from HyDrop v2 data (top) and 10x data (bottom) **e**, Multiscale footprint of the regions shown in c (upper) and d (bottom) in BICCN (10x multi-ome data). **g-j**, Evaluation of motif importance scores (threshold set on 4.25) overlapping regions (present in HyDrop v2 and 10x data) for cell type identity indicated by the BICCN (10x) data-based model and HyDrp v2 data-based model for oligodendrocytes (**g**), L2 3 (**h**), L5 ET (**i**), and L6 IT neurons (**j**), **k-m**, Mouse-specific Tn5 bias from Hu et al., (2025) plotted on cell type-specific DARs and carrot plot showing the regions of accessible chromatin ±0.5 kb around the center of DARs sorted by the highest (blue) to lowest (red) accessibility in for HyDrop v2, HyDrop v1, 10x v2, and 10x v1 data for oligodendrocytes (**k**), microglia (**l**), and L2 3 neurons (**m**).

**Fig. S5.**
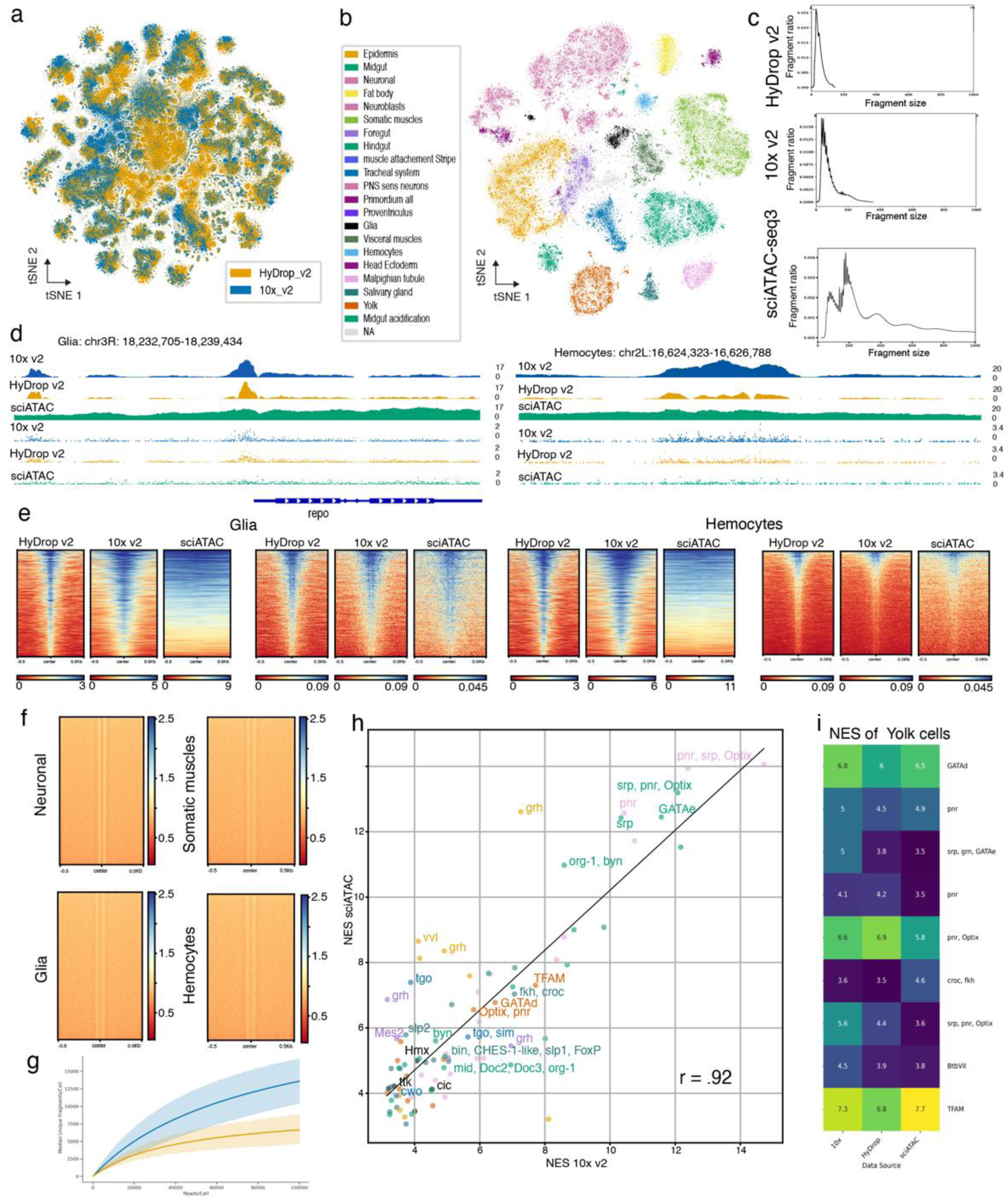
HyDrop v2 captures cellular diversity comparable to 10x and more accurate than sciATAC in developing Drosophila melanogaster. **a,** *t*-distributed stochastic neighbor embedding (tSNE) of all 607,330 cells generated with HyDrop v2 (340,604, 18 experiments) and 10x v2 (266,736 cells, 5 experiments) colored by technique, batch correction according to wet lab protocol used to generate data. Data points were randomly shuffled before plotting. **b,** *t*-distributed stochastic neighbor embedding (tSNE) of 61,480 cells of 16-20h AEL extracted from sciATAC embryo atlas (Calderon et al., 2022) colored by cell type. Data points were randomly shuffled before plotting. **c,** Fragment size plot of HyDrop v2, 10x v2 and sciATAC-seq3 experiments. d, Genome tracks of cell-type specific DARs down sampled to the smallest count present for each technique, 1,247 cells in glia and 1,002 cells in hemocytes. The shown genome tracks are normalized for the fragment count. **d,** Carrot plot showing the regions of accessible chromatin ±0.5 kb around the center of DARs sorted by the highest (blue) to lowest (red) accessibility for glia and hemocytes for HyDrop v2, 10x v2, and sciATAC data. Per cell type, the left plots are generated using the coverage while the right plots are generated using the cut site information extracted from cell-type specific fragment files. **e,** Drosophila-specific Tn5 bias from Hu et al., (2025) plotted on DARs of neuronal cells, somatic muscle cells, glia, and hemocytes. **f,** Sequencing efficiency compared between HyDrop v2 and 10x v2 samples. The colors are shown in a, the labels are shown for the top three motifs per cell type. **g,** Scatterplot of normalized enrichment score of cell-type-specific transcription motifs, dots represent a common motif between sciATAC and 10x v2 data colored per cell type as shown in b. h, Heatmap for top 1000 DARs Yolk cells of 10x v2, HyDrop v2, and sciATAC data. DAR: differentially accessible region.

**Fig. S6.**
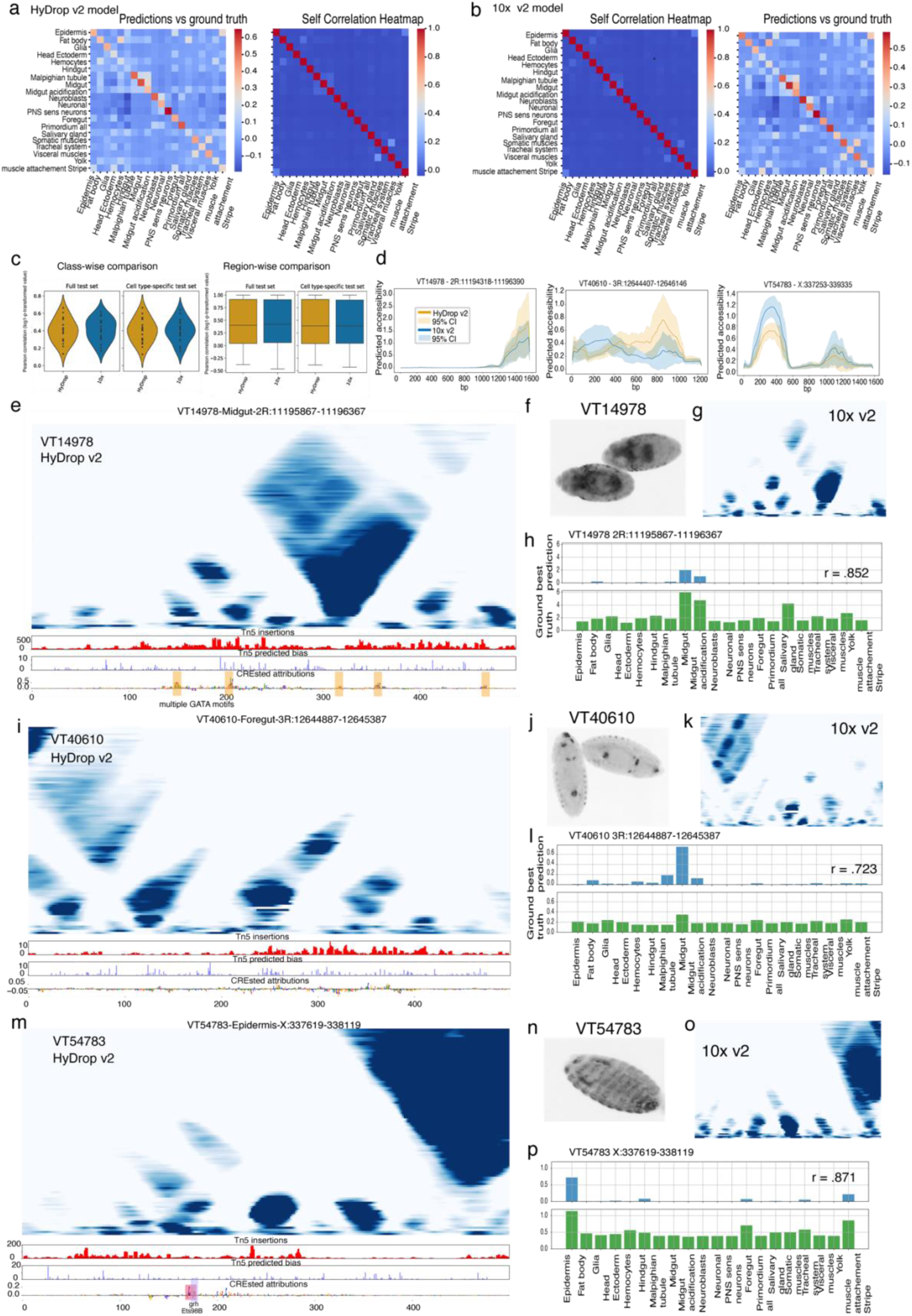
HyDrop v2 sequence model matches results of 10x v2 sequence model and footprinting. **a**, Heatmap of correlation of prediction by HyDrop v2 model with ground truth (left), self-correlation of HyDrop v2 data as ground truth (right) for HyDrop v2 data. **b**, Heatmap of correlation of prediction by 10x v2 model with ground truth (right), self-correlation of 10x v2 data as ground truth (left) for 10x v2 data. **c**, Model comparison of sequence models trained on HyDrop v2 data (orange) and the 10x v2 data set (blue). The accuracy of the predictions evaluated based on the ground truth of region accessibility is compared on class-wise (top) and region-wise level (bottom) for both the full test set in the data and cell type-specific test sets. **d**, Predicted accessibility of 500 bp windows of VT3067 enhancer from VDRC library. Both, the predictions based on the HyDrop v2 and 10x v2 data are shown confidence intervals reflect 10-fold cross-validation models. **e**, Multiscale footprint of 500bp region in VT14978 in midgut cells from HyDrop v2 data only. The bottom track shows Tn5 insertion, the predicted Tn5 bias, and the nucleotide contribution scores based on the HyDrop v2 sequence model. Previously described TF binding sites are highlighted for the corresponding TF. **f,** *In situ* hybridization of VT14978-Gal4 reporter embryos stage 15 with antisense Gal4 probe by Kvon et al. (2014). **g**, Multiscale footprint of 500bp region in VT14978 in midgut cells from 10x v2 data. **h**, Prediction (top) versus ground truth (bottom) of 500bp region within 2kb long VDRC enhancer VT14978 active in midgut cluster. **i**, Multiscale footprint of 500bp region in VT40610 in foregut cells from HyDrop v2 data only. The bottom track shows Tn5 insertion, the predicted Tn5 bias, and the nucleotide contribution scores based on the HyDrop v2 sequence model. Previously described TF binding sites are highlighted for the corresponding TF. **j,** *In situ* hybridization of VT40610-Gal4 reporter embryos stage 15 with antisense Gal4 probe by Kvon et al. (2014). **k**, Multiscale footprint of 500bp region in VT40610 in foregut cells from 10x v2 data. **l**, Prediction (top) versus ground truth (bottom) of 500bp region within 2kb long VDRC enhancer VT40610 active in midgut cluster. **m**, Multiscale footprint of 500bp region in VT54783 in epidermis cells from HyDrop v2 data only. The bottom track shows Tn5 insertion, the predicted Tn5 bias, and the nucleotide contribution scores based on the HyDrop v2 sequence model. Previously described TF binding sites are highlighted for the corresponding TF. **j,** *In situ* hybridization of VT54783-Gal4 reporter embryos stage 15 with antisense Gal4 probe by Kvon et al. (2014). **k**, Multiscale footprint of 500bp region in VT54783 in epidermis cells from 10x v2 data. **l**, Prediction (top) versus ground truth (bottom) of 500bp region within 2kb long VDRC enhancer VT54783 active in midgut cluster. VDRC: Vienna Drosophila Resource Center

**Fig. S7.**
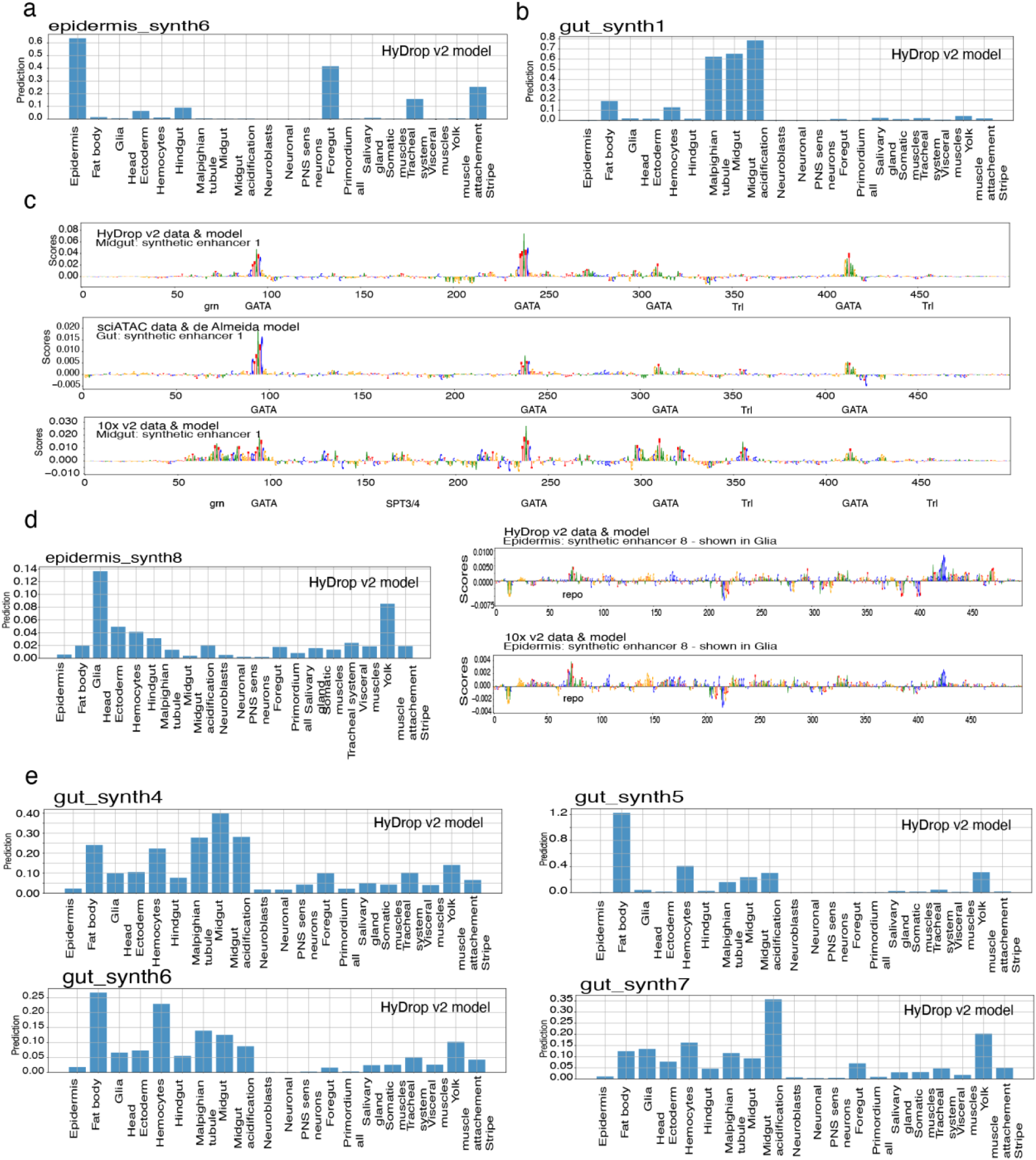
Evaluation of synthetic enhancers designed by de Almeida et al (2023) by HyDrop v2 model. **a**, Prediction of tissue accessibility of synthetic enhancer designed for epidermis cells in 10-12h embryos. Enhancers were designed by de Almeida et al. (2023) and scored with the HyDrop v2 model. Accessibility is specifically predicted for epidermis cells in the HyDrop v2 data. **b**, Prediction of tissue accessibility of synthetic enhancer designed for gut cells in 10-12h embryos. Accessibility is specifically predicted for midgut cells in the HyDrop v2 data. **c**, Nucleotide contribution score of the 500 bp of synthetic enhancer designed to be active in gut cells based on sciATAC data by de Almeida et al. (2023): HyDrop v2 and 10x v2 model predictions shown for midgut cells versus predictions by the original model for general gut cells. **d,** Prediction of tissue accessibility of synthetic enhancer designed for epidermis cells in 10-12h embryos. Accessibility is specifically predicted for glial cells in the HyDrop v2 data (left). Right: Nucleotide contribution score of the 500 bp of synthetic enhancer designed to be active in epidermis cells based on sciATAC data by de Almeida et al. (2023): HyDrop v2 and 10x v2 model predictions shown for glial cells instead showing a *repo* motif. **e**, Prediction of tissue accessibility of synthetic enhancer designed for gut cells in 10-12h embryos. Enhancers were designed by de Almeida et al. (2023) but did not give signal *in vivo*. Here, the synthetic sequences are scored with the HyDrop v2 model showing predicted accessibility in other tissues than the gut.

**Table S1.**
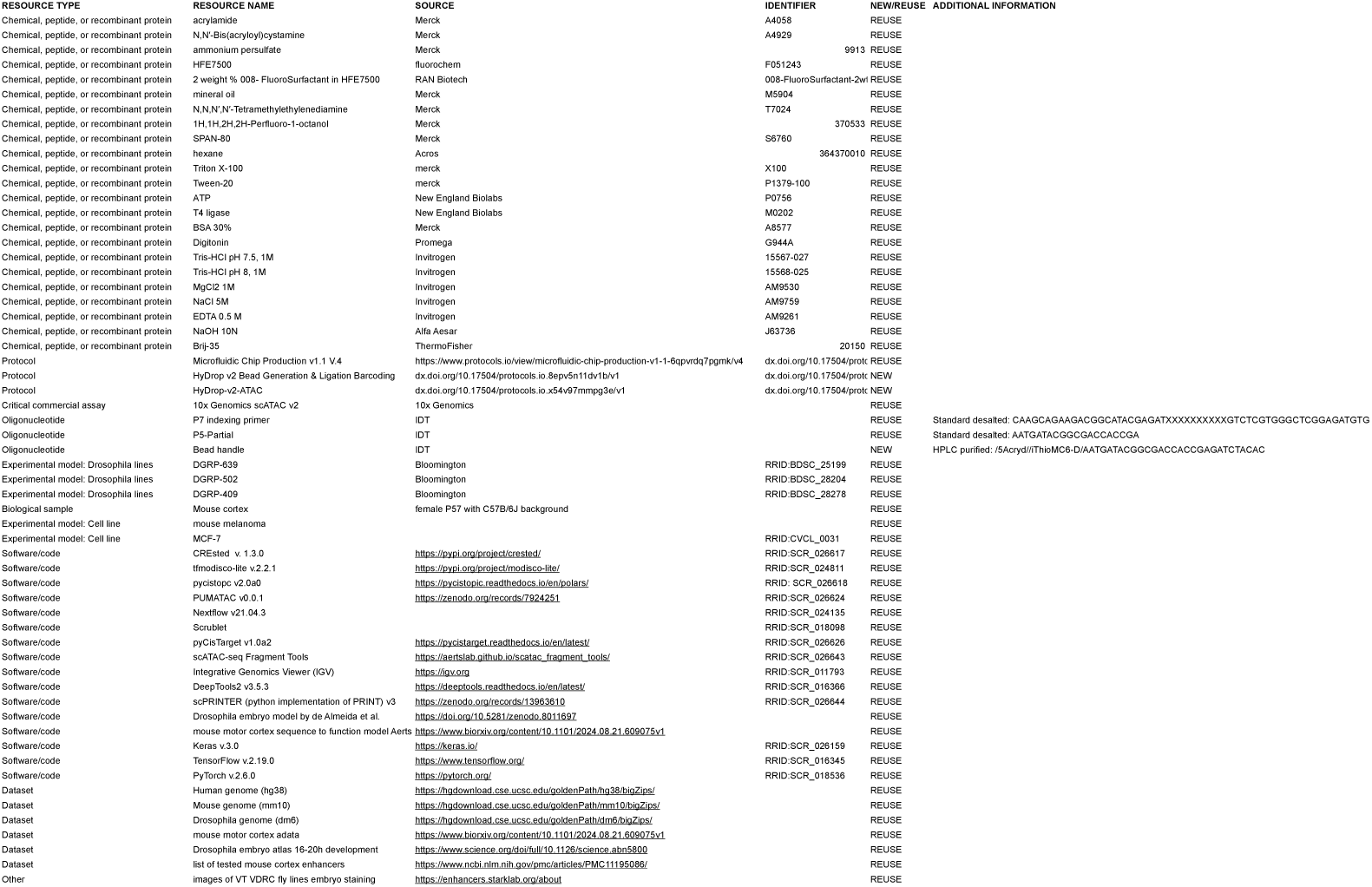
Key resource table.

**Table S2.** VTID enhancers stage 15 and 16 by Kvon et al. (2014) annotated according to HyDrop Drosophila annotation.

| VTID | Dm6 Chromo | Dm6 Start | Dm6 End | stg15_16 | HyDrop_cell_types |
| --- | --- | --- | --- | --- | --- |
| VT1083 | chr2L | 2163674 | 2165773 | antenna-maxillary-labial_complex;3\|brain_subset;3 | Neuronal |
| VT11145 | chr2L | 21843469 | 21845803 | head_subset;3\|dorsal_vessel;3\|midgut_chamber;3 | Midgut, Head_Ectoderm, Midgut_acidification |
| VT14335 | chr2R | 9930287 | 9932397 | NA | NA |
| VT14616 | chr2R | 10459397 | 10461789 | hindgut;5\|anal_pad;4\|atrium;4 | Hindgut |
| VT14978 | chr2R | 11194318 | 11196390 | amnioserosa;5\|midgut_broad;3\|ubiquitous;3 | Midgut |
| VT15726 | chr2R | 12593526 | 12595724 | head_subset;3\|head_epidermis_broad;3\|pharynx;3\|trunk_epidermis | Epidermis, Head_Ectoderm, Foregut, Hindgut |
| VT1597 | chr2L | 3113886 | 3116075 | salivary_gland;5\|gonad;2 | Salivary_gland |
| VT16530 | chr2R | 14173216 | 14175346 | ubiquitous;4 | NA |
| VT16613 | chr2R | 14334875 | 14336968 | antenna-maxillary-labial_complex;3\|amnioserosa;5 | NA |
| VT17409 | chr2R | 15897411 | 15899572 | ventral_nerve_cord_subset;4\|anal_pad;3\|brain_subset;4\|malpighian_tubule | Neuronal, Malpighian_tubule, Head_Ectoderm |
| VT17873 | chr2R | 16783707 | 16785865 | yolk;4\|hemocyte;3\|macrophage;3 | Yolk, Hemocytes |
| VT18684 | chr2R | 18338586 | 18340728 | brain_broad;5\|ventral_nerve_cord_broad;5 | Neuronal, Head_Ectoderm |
| VT19073 | chr2R | 19107715 | 19109845 | head_epidermis_broad;5\|trunk_epidermis_broad;5 | Epidermis |
| VT19429 | chr2R | 19819410 | 19821552 | tracheal_system_subset;3\|anterior_spiracle;3 | Tracheal_system |
| VT19535 | chr2R | 20026362 | 20028504 | head_subset;3\|anal_pad;5 | Neuronal, Head_Ectoderm |
| VT19841 | chr2R | 20597698 | 20599839 | head_subset;2 | Neuronal, Head_Ectoderm |
| VT21015 | chr2R | 22802043 | 22803585 | NA | NA |
| VT21151 | chr2R | 23049881 | 23051973 | NA | NA |
| VT21433 | chr2R | 23592106 | 23594358 | foregut;3 | Proventriculus |
| VT21819 | chr2R | 24327840 | 24330065 | head_subset;4\|midgut_broad;3\|head_epidermis_subset;5\|trunk_epidermis | Neuronal, Midgut, Epidermis, Midgut_acidification |
| VT22068 | chr2R | 24807620 | 24809727 | NA | NA |
| VT22223 | chr2R | 25106786 | 25108894 | telson_subset;4\|head_epidermis_ventral_broad;3 | Epidermis, Head_Ectoderm |
| VT23462 | chr3L | 373888 | 375960 | head_subset;4\|salivary_gland_common_duct;4 | Neural, Salivary_gland |
| VT23835 | chr3L | 1095173 | 1097593 | lateral_epidermis_subset;3\|ventral_midline;3\|midgut_broad;3 | Epidermis, Midgut, Midgut_acidification |
| VT23951 | chr3L | 1322658 | 1324778 | trunk_epidermis_broad;3 | Epidermis |
| VT26739 | chr3L | 6806613 | 6808816 | trunk_epidermis_broad;5 | Epidermis |
| VT28557 | chr3L | 10351940 | 10354125 | head_subset;4\|somatic_muscle;5 | Neural, Somatic_muscles, Head_Ectoderm |
| VT29804 | chr3L | 12693742 | 12696663 | midgut_chamber;2\|proventriculus;4 | Midgut, Proventriculus |
| VT3067 | chr2L | 6073895 | 6076535 | midgut_broad;2\|ventral_nerve_cord_broad;4\|trunk_epidermis | Midgut, Neuronal, Head_Ectoderm, Midgut_acidification, Neuroblasts |
| VT31409 | chr3L | 15763704 | 15765967 | antenna-maxillary-labial_complex;3\|telson_subset;3\|midgut | Midgut, Midgut_acidification |
| VT31552 | chr3L | 16043310 | 16045539 | brain_broad;2 | Neuronal, Neuroblasts |
| VT32972 | chr3L | 18787471 | 18789555 | salivary_gland;5\|fat_body;2 | Salivary_gland, Fat_body |
| VT36347 | chr3R | 4451213 | 4453364 | midgut_broad;3\|ubiquitous;5\|head_epidermis_subset;5\|trunk_epidermis | Midgut, Epidermis, Head_Ectoderm, Midgut_acidification |
| VT38601 | chr3R | 8787084 | 8789298 | head_subset;4\|lateral_epidermis_subset;3\|ventral_epidermis | Epidermis, Head_Ectoderm |
| VT39498 | chr3R | 10513249 | 10515382 | head_subset;2\|amnioserosa;4\|midgut_chamber;4 | Midgut |
| VT39530 | chr3R | 10569672 | 10571928 | brain_broad;4\|ventral_nerve_cord_broad;4 | Neuronal, Neuroblasts |
| VT40599 | chr3R | 12621701 | 12623889 | salivary_gland;5 | Salivary_gland |
| VT40610 | chr3R | 12644407 | 12646146 | pharynx;4\|proventriculus;5\|anterior_hindgut;5\|esophagus;4 | Foregut, Hindgut, Proventriculus |
| VT40611 | chr3R | 12645563 | 12648106 | midgut_chamber;2 | Midgut, Midgut_acidification |
| VT40612 | chr3R | 12648083 | 12650858 | head_subset;3 | Neuronal, Neuroblasts |
| VT41025 | chr3R | 13428001 | 13428951 | NA | NA |
| VT41037 | chr3R | 13448589 | 13450735 | visceral_muscle_broad;4 | Visceral_muscles |
| VT42009 | chr3R | 15307864 | 15310235 | somatic_muscle;5 | Somatic_muscles |
| VT42358 | chr3R | 16005428 | 16007531 | midgut_broad;2\|proventriculus;4\|salivary_gland;4\|malpighia | Midgut, Salivary_gland, Malpighian_tubule, Proventriculus |
| VT4243 | chr2L | 8421920 | 8424099 | antenna-maxillary-labial_complex;5\|ventral_nerve_cord_sub | Neuronal, Neuroblasts, Hemocytes |
| VT42746 | chr3R | 16699060 | 16701215 | salivary_gland;5\|outer_optic_lobe;1 | Salivary_gland, Neuronal |
| VT42747 | chr3R | 16700797 | 16702888 | head_subset;3 | Neuronal |
| VT42831 | chr3R | 16890314 | 16892448 | ventral_nerve_cord_subset;3\|brain_subset;3\|dorsal_epidermis | Neuronal, Neuroblasts, Epidermis |
| VT42855 | chr3R | 16936443 | 16938643 | head_subset;3\|lateral_epidermis_subset;3\|telson_subset;3\|h | Epidermis, Neuronal, Neuroblasts |
| VT42863 | chr3R | 16950676 | 16952842 | pharynx;3\|trunk_epidermis_subset;3 | Foregut, Epidermis |
| VT4320 | chr2L | 8578321 | 8580523 | ventral_epidermis_subset;3\|trunk_epidermis_subset;1\|dorsal | Epidermis, Head_Ectoderm |
| VT43781 | chr3R | 18695700 | 18697823 | pharynx;5\|atrium;3 | Foregut |
| VT44101 | chr3R | 19300581 | 19302962 | pharynx;2\|telson_subset;2\|proventriculus;5 | Foregut, Midgut |
| VT44107 | chr3R | 19314425 | 19316666 | ventral_epidermis_subset;2\|brain_subset;3 | Neuronal, Neuroblasts, Head_Ectoderm |
| VT4448 | chr2L | 8841573 | 8843738 | ventral_midline;2\|brain_subset;3 | Neuronal |
| VT45611 | chr3R | 22168686 | 22170923 | head_subset;3\|lateral_epidermis_subset;3\|telson_broad;5 | Epidermis |
| VT45651 | chr3R | 22253586 | 22255816 | head_subset;5\|stomatogastric_nervous_system;5 | PNS_sens_neurons |
| VT47167 | chr3R | 25171908 | 25174205 | NA | NA |
| VT47408 | chr3R | 25632148 | 25634323 | brain_subset;2 | Neuronal |
| VT47619 | chr3R | 26026378 | 26028970 | head_subset;2\|hypopharynx;3 | Neuronal, PNS_sens_neurons |
| VT47640 | chr3R | 26064120 | 26066252 | salivary_gland;5 | Salivary_gland |
| VT48943 | chr3R | 28588434 | 28590577 | midgut_chamber;2\|brain_broad;1\|hindgut;2\|ventral_nerve_c | Midgut, Neuronal, Head_Ectoderm, Midgut_acidification |
| VT48946 | chr3R | 28592856 | 28595111 | brain_subset;1 | Neuronal |
| VT49445 | chr3R | 29562062 | 29564342 | lateral_epidermis_subset;5 | Epidermis |
| VT49668 | chr3R | 29983451 | 29986358 | head_epidermis_broad;5\|pharynx;5\|trunk_epidermis_broad;3 | Epidermis, Foregut, Head_Ectoderm |
| VT50195 | chr3R | 30923617 | 30925849 | head_subset;3\|midgut_chamber;4\|midgut_interstitial_cell;4 | Midgut |
| VT54783 | chrX | 337253 | 339335 | ventral_epidermis_subset;3\|salivary_gland;5\|head_epidermis | Epidermis, Salivary_gland |
| VT55906 | chrX | 2651914 | 2654026 | head_epidermis_broad;5\|pharynx;4\|trunk_epidermis_broad;3 | Foregut, Epidermis |
| VT57083 | chrX | 5054760 | 5056847 | head_epidermis_broad;5\|trunk_epidermis_broad;5\|ubiquitous | Epidermis |
| VT58723 | chrX | 8354492 | 8356703 | antenna-maxillary-labial_complex;5\|lateral_epidermis_subse | Epidermis |
| VT58999 | chrX | 8882617 | 8884691 | antenna-maxillary-labial_complex;3\|pharynx;5\|proventriculu | Midgut, Foregut, Proventriculus |
| VT59092 | chrX | 9059967 | 9062117 | midgut_chamber;5\|head_epidermis_subset;4\|trunk_epiderm | Midgut, Epidermis, Midgut_acidification, Head_Ectoderm |
| VT62606 | chrX | 16169462 | 16171589 | visceral_muscle_broad;5 | Visceral_muscles |
| VT6436 | chr2L | 12589222 | 12591343 | amnioserosa;3\|midgut_broad;5\|salivary_gland;4 | Midgut, Salivary_gland, Head_Ectoderm |
| VT64887 | chrX | 20651972 | 20654082 | ventral_nerve_cord_subset;1\|brain_subset;1 | Neuronal, Neuroblasts |
| VT7402 | chr2L | 14485084 | 14487223 | head_epidermis_lateral_subset;2 | Epidermis |
| VT7922 | chr2L | 15480181 | 15482932 | NA | NA |
| VT8346 | chr2L | 16317691 | 16319794 | posterior_hindgut;3\|proventriculus;3\|salivary_gland;5 | Hindgut, Proventriculus, Salivary_gland |
| VT9081 | chr2L | 17748587 | 17750842 | NA | NA |
| VT9675 | chr2L | 18890105 | 18892283 | ventral_nerve_cord_subset;2\|amnioserosa;4\|brain_subset;2 | Neuronal, Neuroblasts, Epidermis |

## Notes

### Competing Interest Statement

The authors have declared no competing interest.

### Summary of Updates

Section on Data Availability updated to make sequence-to-function models available.

